# Genomic innovation and horizontal gene transfer shaped plant colonization and biomass degradation strategies of a globally prevalent fungal pathogen

**DOI:** 10.1101/2022.11.10.515791

**Authors:** Neha Sahu, Boris Indic, Johanna Wong-Bajracharya, Zsolt Merényi, Huei-Mien Ke, Steven Ahrendt, Tori-Lee Monk, Sándor Kocsubé, Elodie Drula, Anna Lipzen, Balázs Bálint, Bernard Henrissat, Bill Andreopoulos, Francis M. Martin, Christoffer Bugge Harder, Daniel Rigling, Kathryn L. Ford, Gary D. Foster, Jasmyn Pangilinan, Alexie Papanicolaou, Kerrie Barry, Kurt LaButti, Máté Virágh, Maxim Koriabine, Mi Yan, Robert Riley, Simang Champramary, Krista L. Plett, Igor V. Grigoriev, Isheng Jason Tsai, Jason Slot, György Sipos, Jonathan Plett, László G. Nagy

**Affiliations:** Biological Research Center, Synthetic and Systems Biology Unit, 6726 Szeged, Hungary; Doctoral School of Biology, Faculty of Science and Informatics, University of Szeged, 6726 Szeged, Hungary; Functional Genomics and Bioinformatics Group, Faculty of Forestry, Institute of Forest and Natural Resource Management, University of Sopron, 9400 Sopron, Hungary; Hawkesbury Institute for the Environment, Western Sydney University, Richmond, New South Wales, Australia; Elizabeth Macarthur Agricultural Institute, NSW Department of Primary Industries, Menangle, NSW, 2568, Australia; Biodiversity Research Center, Academia Sinica, Taipei, Taiwan; U.S. Department of Energy Joint Genome Institute, Lawrence Berkeley National Laboratory, Berkeley, CA 94720, USA; Department of Microbiology, Faculty of Science and Informatics, University of Szeged, Szeged, Hungary; ELKH-SZTE Fungal Pathogenicity Mechanisms Research Group, University of Szeged, 6726 Szeged, Hungary; Architecture et Fonction des Macromolécules Biologiques (AFMB), CNRS, Aix-Marseille Université, Marseille, France.; INRAE, UMR 1163, Biodiversité et Biotechnologie Fongiques, Marseille, France.; DTU Bioengineering, Technical University of Denmark, 2800 Kongens Lyngby, Denmark; Department of Biological Sciences, King Abdulaziz University, Jeddah 999088, Saudi Arabia; Université de Lorraine, INRAE, UMR 1136 ‘Interactions Arbres/Microorganismes’, Centre INRAE Grand Est - Nancy, 54280 Champenoux, France; Department of Biology, Section of Terrestrial Ecology, University of Copenhagen, Universitetsparken 23, 2100 København Ø; Department of Biosciences, University of Oslo, PO Box 1066, Blindern, 0316 Oslo, Norway; Swiss Federal Research Institute WSL, 8903 Birmensdorf, Switzerland; School of Biological Sciences, Life Sciences Building, University of Bristol, Bristol BS8 1TQ, UK; Department of Plant and Microbial Biology, University of California Berkeley, Berkeley, CA 94720, USA; Department of Plant Pathology, The Ohio State University, Columbus, Ohio, USA

## Abstract

Members of the fungal genus *Armillaria* are necrotrophic pathogens with efficient plant biomass-degrading strategies. The genus includes some of the largest terrestrial organisms on Earth, spreading underground and causing tremendous losses in diverse ecosystems. Despite their global importance, the mechanism by which *Armillaria* evolved pathogenicity in a clade of dominantly non-pathogenic wood-degraders (Agaricales) remains elusive. Here, using new genomic data, we show that *Armillaria* species, in addition to widespread gene duplications and *de novo* gene origins, appear to have at least 775 genes that were acquired via 101 horizontal gene transfer (HGT) events, primarily from Ascomycota. Functional and expression data suggest that HGT might have affected plant biomass-degrading and virulence abilities of *Armillaria*, two pivotal traits in their lifestyle. We further assayed gene expression during root and cambium colonization, and report putative virulence factors, extensive regulation of horizontally acquired and wood-decay related genes as well as novel pathogenicity-induced small secreted proteins (PiSSPs). Two PiSSPs induced necrosis in live plants, suggesting they are potential virulence effectors conserved across *Armillaria*. Overall, this study details how evolution knitted together horizontally and vertically inherited genes in complex adaptive traits, such as plant biomass degradation and pathogenicity, paving the way for development of infection models for one of the most influential pathogens of temperate forest ecosystems.

## Introduction

Plant pathogenic fungi cause significant economic losses worldwide in a wide variety of plant species, including forest trees. Among tree pathogens, the genus *Armillaria* (Basidiomycota, Agaricales, Physalacriaceae) stands out as one of the most important in temperate systems, responsible for great losses in both natural and planted stands of woody plants ^1–3^. Known at the genus-level as *Armillaria* root-rot disease ^3–5^, the most common pathogenic species is *A. mellea sensu lato,* which has been reported to infect >500 plant species ^1, 6, 7^ and is solely responsible for up to 40% annual loss of vinegrape in California ^8^. *A. ostoyae* is responsible for significant losses in conifer forests ^9, 10^.

*Armillaria* species evolved a range of features exceptional among fungi, which have conceivably all emerged in its most recent common ancestor (MRCA). These include a very low mutation rate, extreme longevity and immense colony sizes (>2,500 years, >900 hectares ^1, 3, 11– 13^), diploidy ^14^, bioluminescence ^15^, specialized underground structures known as rhizomorphs ^16, 17^, and the potential to fix atmospheric N_2_ ^18^. Perhaps the most economically important aspect of this genus is the ability to infect and kill woody plants ^1, 2, 19, 20^. Most *Armillaria* species are broad host range necrotrophs ^1, 3^. After infection through the roots, they colonize and kill the cambium of the tree, causing the death of the plant and enabling the fungus to transition to its necrotrophic phase ^1, 21^. Despite a well-documented epidemiology and etiology ^2, 19, 21–24^, molecular aspects of the infection process are poorly known. Recent studies and genome sequencing efforts highlighted certain secondary metabolites, plant cell wall degrading enzymes (PCWDEs), chitin-binding proteins and expanded protein-coding repertoires enriched in putative pathogenicity-related genes, among others ^1, 17, 18, 25–30^. It is likely that these previous findings only cover a fraction of the virulence genes of *Armillaria*, leaving much of its pathogenic arsenal yet to be characterized.

As a result, it is unknown how infection models developed based on better-studied necrotrophic fungi (e.g. *Sclerotinia sclerotiorum* ^31–33^, or *Colletotrichum* spp. ^34^), are applicable for *Armillaria*, if at all, and what traits the latter evolved for plant infection, are unknown. In other necrotrophs, broad roles of tissue acidification, tolerance towards and detoxification of plant secondary metabolites and reactive oxygen bursts ^35–37^, secretion of diverse plant cell wall degrading enzymes (PCWDEs) ^34, 38^ and effectors ^39^ have been established as key assets of infection. However, *Armillaria* represents an independent origin of necrotrophy, which might have resulted in unique infection mechanisms, our understanding of which remains limited.

Here, we sequenced eight new genomes and report transcriptome data from new *in planta* and *in vitro* pathosystems, allowing us to explore genome evolution and key aspects of the necrotrophic lifestyle of *Armillaria* spp. We infer that gene duplications, genus-specific gene families and horizontal gene transfers have shaped the genomic toolkit available for plant infection and biomass degradation. RNA-Seq data from six experiments and four *Armillaria* species, including new *in planta* time series- and fresh stem invasion experiments allowed us to decipher gene expression patterns specific for these processes in virulent and non-virulent strains. Experimental validation of predicted pathogenicity-induced small secreted proteins revealed potential conserved virulence factors in *Armillaria*. Overall, our phylogenomic and gene expression studies elucidate how vertically and horizontally acquired genes became integrated into complex adaptive traits of a globally important fungal group.

## Results

### New *Armillaria* genomes

We report the high-quality annotated *de novo* genomes of eight *Armillaria* species including *A. borealis*, *A. ectypa*, *A. fumosa*, *A. luteobubalina, A. mellea*, *A. nabsnona*, *A. novae-zealandiae* and *A. tabescens*. The new genomes were assembled with Falcon ^40^ to 33-864 scaffolds comprising 40-79 Mbp haploid size, with 12,228 - 19,984 predicted gene models and BUSCO (fungi) completeness 97.7-99.7% (Figure 1). Sequencing statistics for the new genomes are summarized in Table S1.

**Figure 1:**
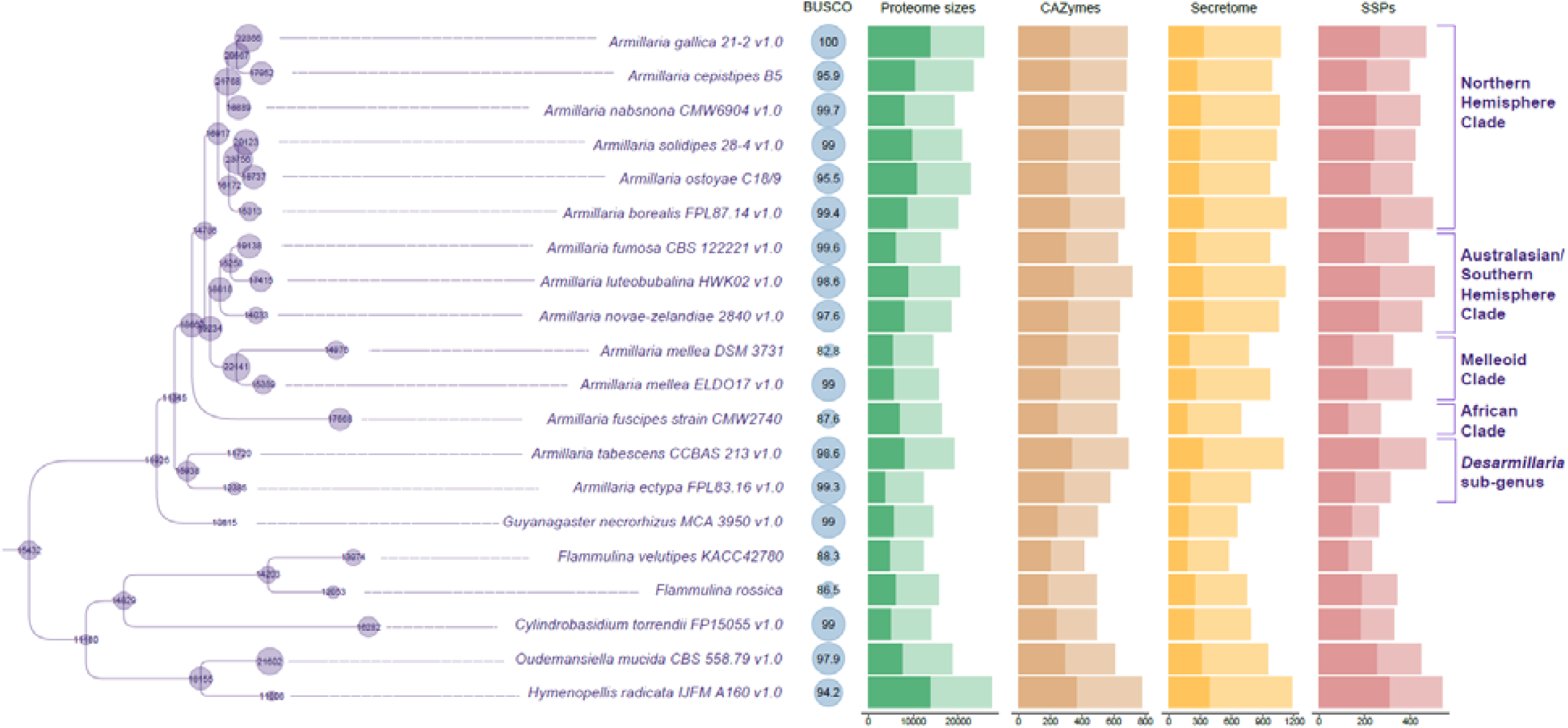
General genome statistics and reconstruction of ancestral genome size dynamics for 15 *Armillaria* species and 5 Physalacriaceae outgroups. Numbers at nodes represent ancestral proteome sizes in the Physalacriaceae tree. Purple dot corresponds to sizes for each node in the tree (for gene gains and losses at each node, and for the complete species tree, see Figure S1). Blue dots represent BUSCO scores for each species. For proteome sizes, darker color shows proteins with no known functional domains (unannotated proteins). Carbohydrate active enzymes (CAZymes) - darker color shows plant cell wall degrading enzymes (PCWDEs), lighter color shows other CAZymes. Secretomes and SSPs - darker color shows unannotated proteins, lighter color shows proteins with known functional domains (for general genome statistics of all 66 species in Dataset 1, see Table S1).

We sampled all major clades recognized currently ^41^, including the Northern Hemisphere (*A. borealis, A. ostoyae, A. gallica, A. cepistipes, A. solidipes, A. nabsnona*), the Australasian/Southern American (*A. luteobubalina, A. fumosa, A. novae-zelandiae*), African (*A. fuscipes*) and melleoid clades (*A. mellea*). We also included 2 species from subgenus *Desarmillaria* (*A. tabescens* and *A. ectypa*), of which the moss-associated *A. ectypa* had the smallest genome among *Armillaria* spp., as well as *Guyanagaster necrorhizus* ^18^, which is the sister genus of *Armillaria* ^41^ (Figure 1). For comparative analyses, we annotated carbohydrate active enzymes (CAZymes), plant cell wall degrading enzymes (PCWDEs), secreted proteins and small secreted proteins (SSPs) (<300 amino acids).

### Genetic innovations in *Armillaria* clade

To analyze the genomic innovations associated with the rise of *Armillaria*, we used genomes of 15 *Armillaria* species and 5 outgroups from the Physalacriaceae, which increased sampling density nearly four-fold compared to previous comparative studies ^17, 25^. We combined these with those of other Agaricales exhibiting a range of lifestyles, resulting in a dataset of 66 species (Table S1).

Reconstruction of genome-wide gene gain/loss patterns revealed genome expansion in *Armillaria*. The estimated ancient gene set at the root node comprised 9,929 genes, suggesting an early origin for most genes (Figure S1). The *Armillaria* genus showed a net genome expansion with 18,662 protein-coding genes inferred for the most recent common ancestor (MRCA) of *Armillaria* (2,913 duplications, 189 losses), as opposed to 15,938 for the MRCA of *Armillaria* and *Guyanagaster* (2,158 duplications, 4,375 losses) and 18,155 for the MRCA of *Armillaria*, *Guyanagaster* and *Hymenopellis* (653 duplications, 5,187 losses) (Figure 1, Figure S1). The MRCA of the Northern Hemisphere clade was inferred to have undergone further expansion to 23,736 genes (3,140 duplications, 1,525 losses). These data indicate that the large protein coding repertoires of *Armillaria* spp. can be explained by genus-specific gene duplications, as suggested before ^17^.

The 2,913 duplications in the MRCA of *Armillaria* happened in a total of 1,473 orthogroups (OGs). Gene ontology (GO) and InterPro enrichment analyses for these showed significant overrepresentation of 55 molecular functions, 18 biological processes and 3 cellular component terms (using topGO ^42^, weight01 algorithm, p <0.05) (Figure S2) and 733 InterPro terms (Table S2). These included functions related to plant biomass utilization, such as pectin-degradation (pectate lyases, pectin lyases, pectin esterases, GH28), cellulose binding, and putative extracellular and aromatic compound breakdown (e.g. intradiol ring-cleavage dioxygenase, multicopper oxidases) (see Table S2, Figure S2). Duplicated genes were also enriched in putative pathogenesis related gene families. For instance, we found significant overrepresentation of deuterolysins, aspartic peptidases, chitin deacetylases, and SCP-like Golgi-associated pathogenesis-related proteins. Ceratoplatanins and LysM domains, which were reported to assist infection in pathogenic fungi ^43–45^ were also enriched in *Armillaria*.

Small secreted proteins (SSPs), which include cysteine-rich proteins involved in, among others, host colonization ^46^, were found in 270-507 copies in *Armillaria*, with *A. fuscipes* having the fewest and *A. luteobubalina* having the most. Of these, 45-57% had no known functional domains, which we hereafter refer to as unannotated SSPs. Typically this latter class of SSPs are called candidate secreted effector proteins (CSEPs) in pathogenic fungi ^34, 47, 48^.

Gene families that arose within and are conserved in most *Armillaria* spp., referred to as novel core orthogroups, may be particularly relevant for explaining the plethora of *Armillaria*-specific innovations. We found 212 novel core orthogroups shared by at least twelve *Armillaria* species including *G. necrorhizus* (Table S2, Figure S3). Of these, 116 consisted of proteins with no known functional annotations. The remaining orthogroups were dominated by F-box domains, Leucine-rich repeats, Cytochrome P450s, Zinc-finger C2H2 type transcription factors, protein kinases and other fast evolving gene families (Table S2).

*Armillaria* spp. are among the few bioluminescent fungi ^15^. Ke et al. reported the luciferase gene cluster in *Armillaria,* comprising five genes ^49^. We found the luciferase cluster to be conserved and highly syntenic in all *Armillaria* genomes (Figure S4), however, the cluster was missing in other Physalacriaceae, suggesting it was lost in those, or gained in the common ancestor of *Armillaria* and *Guyanagaster*.

#### Plant biomass degrading enzymes in *Armillaria* and the Physalacriaceae

Ecologically, *Armillaria* species are reported to be facultative necrotrophs that first kill the host, then utilize its biomass during the saprotrophic phase. The new genomes allowed us to make predictions on the wood decay strategy of *Armillaria*, relative to other wood-decaying fungi based on their plant cell wall degrading enzyme genes (PCWDEs). Similar to white rot (WR) fungi and necrotrophs ^39, 50^, *Armillaria* species possess the complete enzymatic repertoire for degrading woody plant biomass (Figure 1). We generated phylogenetic PCAs for *Armillaria* species, other Physalacriaceae, WR and litter decomposer (LD) species (Table S1) based on PCWDEs acting on cellulose, hemicellulose, pectin and lignin. Cellulases and pectinases clearly separated *Armillaria* spp. and other Physalacriaceae from WR and LD fungi (Figure 2, Figure S5), whereas hemicellulase and ligninase PCA-s grouped them together (Figure 2, Figure S6, Figure S7, Figure S8). Cellulase loading factors indicate that this separation was mainly driven by expansins, the AA16, AA8, and AA3_1, GH1 and GH45 families. In line with loading factors, *Armillaria* and other Physalacriaceae have more AA3_1, GH1 and GH45 genes than WR and LD fungi (Figure S9). On the other hand, they were depleted in CBM1 and GH5_5 with an average of 12 and 5 genes respectively, compared to WR and LD genomes encoding 20-60 CBM1 and 4-10 GH5_5 genes. The GH44 family, which is specific to Basidiomycota ^51^, is missing in the Physalacriaceae and *Armillaria*, indicating that gene losses drove trophic mode evolution of these fungi towards an Ascomycota-like lifestyle.

**Figure 2:**
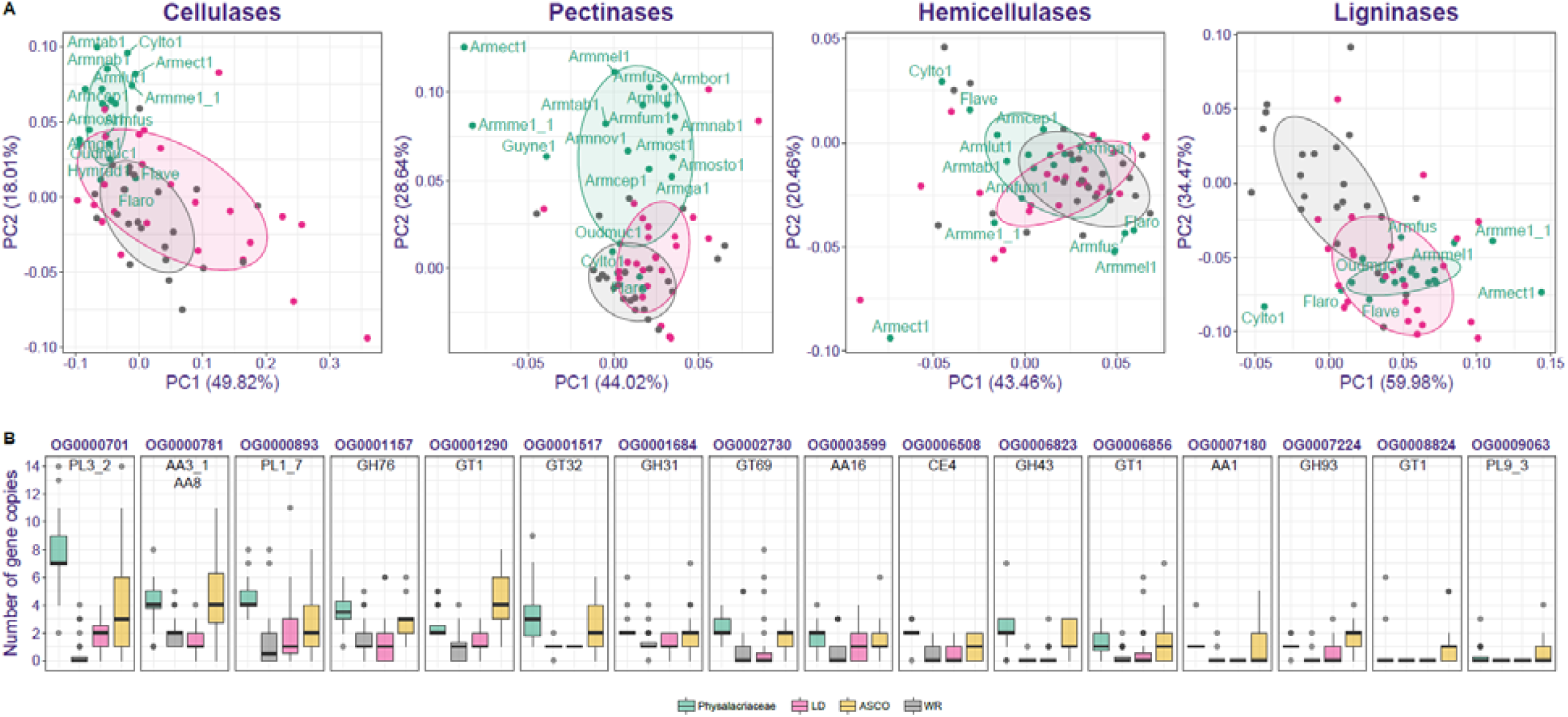
Plant biomass degradation genes in *Armillaria*. A) Phylogenetic PCAs for PCWDE gene families. Species abbreviations are shown only for Physalacriaceae species (all species names and PCA loadings are given in Figure S5 and Table S3). B) Boxplot of copy numbers of 16 CAZy orthogroups co-enriched in Physalacriaceae and in Ascomycota with respect to WR and LD fungi. Scale limits for the boxplot were set to 14, losing one sample point (Conioc1 in OG0000781 with 18 genes). Lifestyles of the species used are denoted by colors for both A and B.

In the pectinase PCA, the highest loading family was CBM67 (rhamnose binding modules), which were enriched in *Armillaria* species and were absent in most WR species in our dataset (Figure S10). CBM67s are frequently associated with GH78 and PL1 proteins ^52^. The PL1 family is present in all Physalacriaceae species, but is depleted or missing in many WR and LD fungi (Figure S10). Other pectin-acting families, such as GH28, GH53, GH88, CE8, and CE12, were present in higher numbers in *Armillaria* and other Physalacriaceae than in WR and LD fungi, whereas PL1_7, PL26, PL3_2, and PL4_1 were abundant in *Armillaria* species but absent in most WR and some LD species.

These analyses portray *Armillaria* and the Physalacriaceae as versatile wood decayers that are nevertheless distinct from WR, despite previous classifications as such ^3, 17, 26, 53, 54^. This is consistent with microscopy, chemical and transcriptomic analyses ^54–59^, which indicated that their decay chemistry is similar to soft rot ^55, 59^, which is known only in the Ascomycota. These observations prompted us to systematically look for similarities between PCWDE repertoires of the Physalacriaceae and the Ascomycota. We found 16 CAZy orthogroups that were significantly overrepresented in both groups with respect to WR+LD fungi (BH-corrected p-value >0.05, Fisher’s exact test (Table S3, Figure 2). These included several of the high-loading families from the PCAs, as well as other PCWDEs acting on cellulose (AA3_1, AA8, CBM1), pectin (PL3_2, PL1_7, PL9_3), cellulose/chitin (AA16), and hemicellulose (GH31, GH43, GH93, CE4). These families could either be the result of co-expansion in both the Ascomycota and the Physalacriaceae or represent horizontal gene transfer (HGT) events. Blast searches with *Armillaria* GH28 genes suggested the latter scenario to be more likely, which led us to systematically evaluate the role of HGTs.

### Widespread horizontal transfer of genes from Ascomycota

To identify horizontally transferred (HT) gene candidates, we calculated the Alien index (AI) ^60^ for each *Armillaria* and Physalacriaceae protein based on a phylogenetically broad set of 942 species from the Ascomycota, Mucoromycota, early-diverging fungi as well as plants and bacteria (Table S1, Dataset3). Based on AI>1, we identified 99-195 HT gene candidates per species, with *A. gallica* having the most and *A. ectypa* the fewest in *Armillaria*. Among Physalacriaceae, *H. radicata* had the highest estimated number of HT genes (247), and *F. velutipes* the lowest (88) (Figure 3A, Table S4).

**Figure 3.**
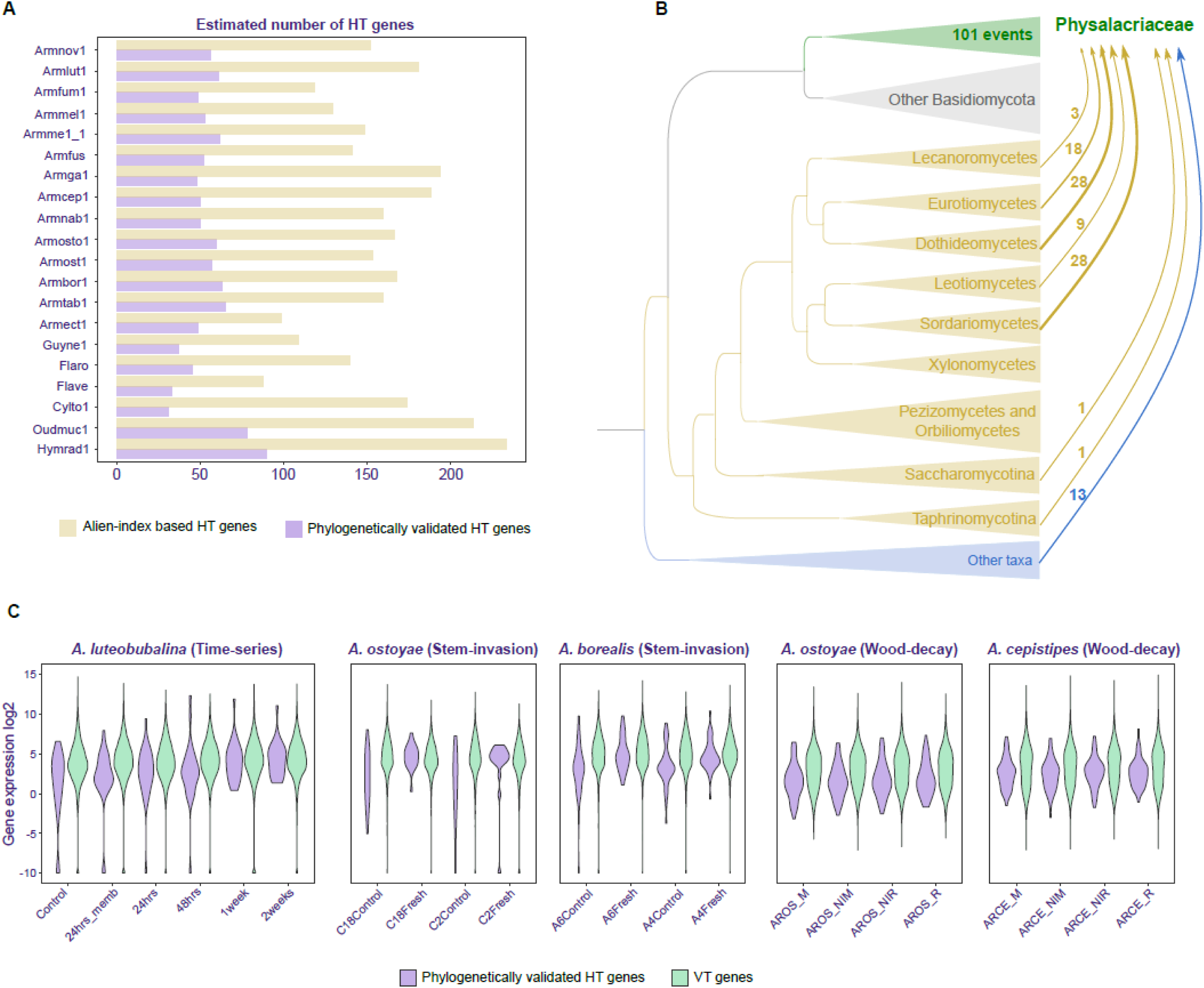
Horizontal gene transfer into *Armillaria* and the Physalacriaceae family. A) Barplot showing the number of HT genes identified in each genome using Alien Index calculations (yellow) and by phylogenetic validation (purple). B) Schematic representation of donor and recipient relationships for HT events after phylogenetic validation. Size of the arrows are proportional to the number of transfer events inferred from all the nodes belonging to a specific donor group. The number of events for each donor group are listed along the arrows. C) Violin plot showing expression dynamics of phylogenetically validated HT genes and vertically transferred (VT) genes. X-axis shows the sample names and y-axis shows the log_2_ transformed gene expression (For expression dynamics of HT and VT genes in developmental dataset, see Figure S11).

Based on the top putative donor species from the AI data, we assembled a dataset comprising the 20 Physalacriaceae, 90 Basidiomycota, 210 Ascomycota, 7 Mucoromycota, and 2 Zoopagomycota species. Using OrthoFinder (v.2.5.4), we identified 649 orthogroups containing at least one gene predicted as HT by AI. Due to the similarity of the CAZy repertoires of Physalacriaceae to Ascomycota (see above), we also included all CAZy orthogroups in these analyses. In total, we analyzed 675 orthogroups by combining phylogenetic support (at least 70% bootstrap in ML gene trees), and top hits against the UniRef100 database ^61^. Overall we recovered 101 strongly supported horizontal transfer events (Figure 3 A) into Physalacriaceae, corresponding to 1089 individual genes. These events were associated with 85 orthogroups, consistent with estimates based on the Alien Index, and an additional 3 CAZy orthogroups, suggesting AI to be an effective strategy for screening HGT candidates. Multiple internal nodes of the Physalacriaceae tree were identified as putative recipients, most of which were deep in the family. 88 HGT events were confidently associated with Ascomycota as donors, in particular the Sordariomycetes and Dothideomycetes (28 transfers each). We find that ∼45% of HT genes have undergone one or multiple rounds of duplication (total 488 duplicated genes) within the Physalacriaceae, indicating that HT genes were likely integrated into the life history of Physalacriaceae and *Armillaria* therein. Expression levels of HT genes support their functionality (Figure S11/Figure 3 C).

Among proteins descended from phylogenetically validated HT events, we found 164 CAZymes (Table S4), of which 117 belonged to families that were co-enriched in Ascomycota and Physalacriaceae compared to WR and LD species (AA3_1, AA8, GH43, GH93, GT1, PL3_2 and PL9_3). This indicates that the co-enrichment signal detected using the simple copy-number based analyses above was likely created by HGT events. In addition to CAZymes, HT genes included intradiol ring-cleavage dioxygenases, CAP domain proteins, Pyr1-like SCP domains, as well as gene families with broad functional roles such as cytochrome P450, peptidases, transporters and transcription factors (Table S4).

Taken together, we find that horizontal transfer affected several families associated with wood-decay (e.g. AA3_1, GH43, PL3_2), as well as plant-fungal interactions (e.g. CAP domain proteins, peptidases), which suggests that it might have shaped the plant biomass degrading and pathogenic abilities of the genus. Given that Physalacriaceae species have been reported to cause soft-rot-like decay on wood ^54–56, 58, 59^ and that soft-rot is classically restricted to the Ascomycota ^62^, we hypothesize that HGT contributed to the evolution of plant biomass degrading ability of these species. We speculate that the large number of putative HGT events from Sordario- and Dothideomycetes might stem from the extensive contact of *Armillaria* ssp. with other plant pathogens in these classes and/or their peculiar niche and longevity. Their rhizomorphs can reach several meters in length and can contact numerous other soil microbes, perhaps providing time windows for gene exchange. Additionally, other idiosyncrasies of *Armillaria*, such as diploidy or the - as yet uncovered - mechanisms that ensure a low mutation rate ^12^, could also be factors in the successful incorporation of foreign DNA in their genomes.

#### *Armillaria* spp. display distinct expression profiles for plant infection, necrotrophy and wood decay

During infection, *Armillaria* enters the host through the root system, colonizes and kills cambium cells, which leads to death of the plant and the onset of the necrotrophic phase ^1^. Molecular aspects associated to this process are hardly known, with most information available on the wood-decaying phase ^17, 58^. To obtain a molecular perspective on these strategies and better understand how *Armillaria* spp. utilize expanding, novel core and HT genes, PCWDEs and gene groups frequently associated with pathogenicity, we produced new RNA-Seq data for two *in vitro* pathosystems (for *A. ostoyae, A. borealis and A. luteobubalina*) and re-analyzed data from 2 published studies ^17, 58^. New data were generated for an *in planta* time-series experiment of *A. luteobubalina* infecting *Eucalyptus grandis* seedlings from 24hrs to 2 weeks (Figure S12 A) and *in vitro* fresh stem invasion assays with highly and less-virulent isolates of both *A. ostoyae* and *A. borealis.* The latter experiment emulated the cambium-killing phase of the fungus (Figure S12 B). Published studies covered wood-decay in *A. ostoyae* and *A. cepistipes* ^58^ as well as rhizomorph and fruiting body development in *A. ostoyae* ^17^.

To obtain a species-independent picture, we aggregated differential gene expression data across experiments and calculated DEG enrichment ratios in each of 24 gene groups we defined (see Methods). These enrichment ratios reflect the proportion of differentially expressed genes in a given gene group in a given experiment and are shown as a heatmap on Figure 4 (see Figure S13 for fruiting body data). *In planta* infection, wood-decay and stem colonization as well as fruiting body/rhizomorph development showed distinct enrichment patterns. Cellulose-, hemicellulose-, pectin- and lignin-related PCWDE genes showed a clear enrichment in stem invasion and wood-decay experiments, as expected based on known functions of these genes ^17, 54^. Among genes upregulated on fresh stems, pectinases were most dominant, which possibly enables the fungus to spread between the bark and the sapwood. At the same time PCWDEs were depleted in the *in planta* time series experiment, indicating that *A. luteobubalina* did not induce these genes during the infection phase. This is consistent with most necrotrophic fungi expressing a limited set of PCWDEs for plant penetration and a larger battery of enzymes during the necrotrophic phase ^34, 38^. Other virulence-related genes include cerato-platanins ^43, 63^, which showed an enriched upregulation at 24h and 48h in *A. luteobubalina* and in stem invasion by *A. borealis* (but not *A. ostoyae*). HT genes, including intradiol ring-cleavage dioxygenases, PCWDEs, peptidases and transporters, showed an enrichment in wood-decay and stem-invasion experiments. Bioluminescence genes were enriched in root-invading mycelium and rhizomorph (*A. ostoyae*) samples. In the developmental dataset SSPs, expanded, novel core and stress-related genes were found to be enriched in various developmental stages of the fungus (Figure S13). Genes related to oxidative stress were upregulated in various stages of *in planta* infection and in stem invasion assays (Figure 4, Figure S14). Superoxide dismutases, catalases and members of the glutathione system and ergothioneine pathway were enriched among upregulated genes in the *in planta* invasion assays, whereas glutathione-S-transferases and catalases were enriched in wood-decay related experiments.

**Figure 4:**
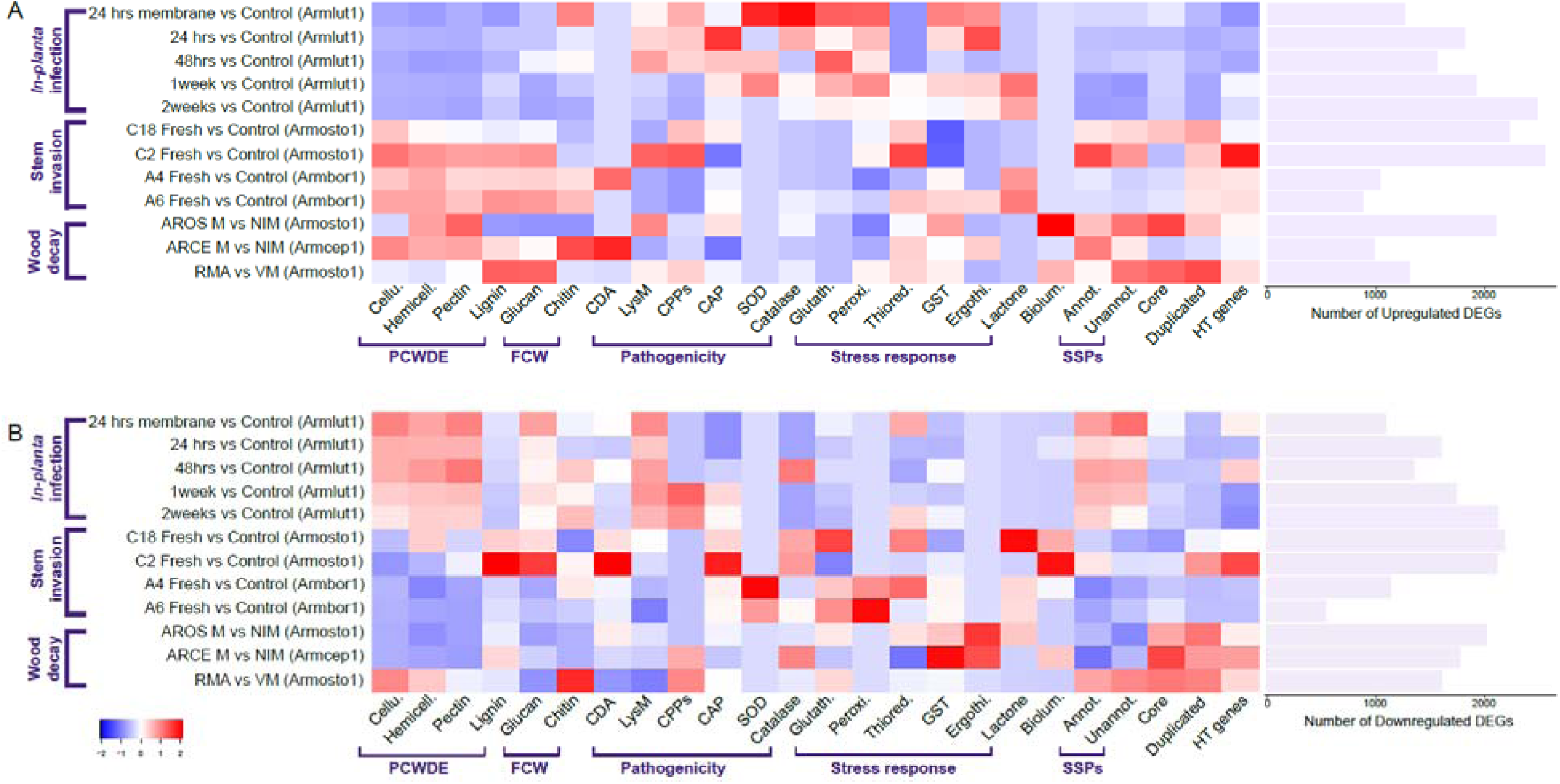
Enrichment of differentially expressed genes of wood-decay, pathogenicity, stress-response and other gene families in 6 RNA-Seq datasets. The heatmap shows enrichment ratios for 24 gene groups (x-axis) from aggregated differential gene expression data across 6 experiments (A - upregulated, B - downregulated genes). Y-axis shows the sample comparison for each dataset, with number of DEGs shown as a barplot at right. In the hatmaps, warmer colors mean higher enrichment ratios (for a complete list of odds ratios Table S5).

It is also evident from Figure 4, that there are species- and strain-specific enrichment patterns among up- and downregulated genes. For example, the virulent *A. borealis* A6 strain had a unique enrichment of pathogenicity-related LysM proteins and cerato-platanins, which was not seen in the less-virulent A4 strain or in *A. ostoyae*. Nevertheless, in both *A. borealis* and *A. ostoyae*, the virulent strains (A6 and C18, respectively) showed more enrichment across most gene categories tested. The species- and strain-specific enrichment patterns also suggest that each species mount a different response under the examined conditions, which might correlate with differences in lifestyle or virulence.

#### Early stages of live host colonization by *Armillaria* characterized by complex transcriptomic regulation

To gain a mechanistic understanding of early stages of *Armillaria* infection, we analyzed gene expression of *A. luteobubalina* colonizing *E. grandis* roots across five time points (pre-symbiosis, 24hrs, 48hrs, 1 week and 2 weeks) in four biological replicates and compared these to non-symbiotic samples (Figure S15, Table S5). SuperSeq revealed that the analysis detected 77-94% of differentially expressed genes (DEGs) in *A. luteobubalina* across the different timepoints, whereas, in *E. grandis*, we could variably detect 25-70% of DEGs (Figure S16). These genes were arranged into modules based on expression similarity using the short time-series expression miner (STEM) ^64^. This yielded eight and four significant modules in the fungus and plant, respectively. Five fungal modules showed gradual increase in expression during time, we refer to these as pathogenicity-specific profiles (Figure S17). Genes in these were enriched for GO terms often associated with pathogenic interactions ^65–67^, with the strongest signal for (oxidative) stress (Figure S17, Figure S14). Inspection of *A. luteobubalina* DEGs identified genes that relate to four pathogenic processes: host immune suppression, oxidative stress, detoxification, and cytotoxicity (e.g. cerato-platanins) ^68^. Glutathione-S-transferases were almost uniformly upregulated in plant-associated samples, compared to free-living mycelium, whereas other genes related to oxidative stress response (e.g. superoxide dismutases) were mostly upregulated at 1 and 2 weeks (Figure S14). Toxin efflux systems, which are involved in the tolerance against plant secondary and defense metabolites (e.g. phytoalexins, phytoanticipins), have been reported to be transcriptionally regulated in necrotrophs ^37^. Detoxification of plant antifungal isoflavones by *A. mellea* has been described ^69^, although the cellular pathways involved are unknown. Membrane transporters, cytochrome p450 monooxygenases and laccases, which are often associated with detoxification or plant metabolites ^37, 70^, showed extensive regulation during the time course of the experiment, with clusters of upregulated genes at 1 and 2 weeks (Figure S14). On the other hand, we did not observe extensive regulation of PCWDEs, which is consistent with patterns reported for data on early colonization of other fungal pathogens (Figure 4).

We observed an upregulation of HT genes at certain stages of *in planta* colonization. Of the 61 phylogenetically validated HT genes in *A. luteobubalina*, 23 were differentially expressed at some time point, however, without any significant enrichment at any individual time point (Figure 4, Table S5). We identified two horizontally acquired CAP domain/pathogenicity-related-1 (PR-proteins which were strongly upregulated across the time-series and three LysM domain proteins (CBM50) that were upregulated at 24 and 48 hours. Both PR-1 and LysM genes are known to be involved in fungal pathogenicity, by facilitating the transport of fatty acids and sterols ^71, 72^ and masking the presence of chitin residues from the plant immune system, respectively ^73, 74^.

When compared to *A. luteobubalina,* DEGs of its host, *E. grandis,* were found to assemble into four significant expression modules. These were enriched in defense-related terms, including chitinase activity, chitin binding, defense response to fungus, defense response to bacterium (Figure S18), suggesting an activation of plant immunity during the early stages of fungal colonization. A finer level analysis of the plant response identified that very few plant genes were differentially regulated across the whole time course, with 95 DEGs up-regulated across the whole time course and 59 DEGs repressed in the same series (Figure S19). Within the core up-regulated genes we found enrichment for phosphoprotein phosphatase activity due to up-regulation of PP2 family genes related to effector immunity and ABA-JA cross-talk ^75^. Other GO terms enriched in this gene set included unfolded protein binding/chaperone activity, induction of heat shock proteins, and jasmonic acid response due to PR4 and MYB108 gene induction ^76, 77^. Of the core repressed genes, we found GO enrichment for copper ion binding proteins, peroxidase activity, carboxylic acid binding, hydrolase activity, and lipid transfer. Reduction of copper ion binding proteins, peroxidase activity and lipid transfer would, likely, lead to a delayed hypersensitive response as copper ions are necessary for ethylene production and repression of the ABA biosynthesis ^78^ while peroxide activity directly leads to plant cell death and the lipid-associated genes encode DIR-like proteins responsible for long-range immune signaling ^79^. The genes associated to carboxylic acid binding and hydrolase activity, meanwhile, likely affect pathogen nutrition more directly as the repressed genes associated to the former are involved in biosynthesis of L-serine, lack of which would reduce the nutrition value of *E. grandis* tissues, while the latter are involved in detoxification (i.e. cyanoalanine nitrilase) which could lead to a toxic build-up of 3-cyano-L-alanine thereby inhibiting pathogen growth ^80^.

#### Putative necrotrophic SSPs identified and tested experimentally

Phytopathogenic fungi often utilize effectors that suppress or manipulate host defense responses ^39, 81, 82^. As many effectors are SSPs, we scrutinized the 39 and 38 annotated and unannotated SSPs, respectively, of *A. luteobubalina* that were upregulated in at least one stage during the time series (Figure 5 and Figure S20). Annotated SSPs with highest fold changes (hereafter FC) included Pry1-like, SCP- or CAP-, Ricin-B-like lectin- and LysM domain containing genes (Table S6), which are related to fungal virulence. Two unannotated SSPs (1165297 and 1348401) had the highest FC values among all genes, so we selected these for experimental validation and call them pathogenicity-induced SSPs (PiSSPs, Figure 5 A, red arrows). Gene 1348401 had a peak expression at 48hrs (FC = 232) whereas 1165297 at 1 week (FC = 447). The encoded proteins contain no conserved domain and have no predicted function. Orthology searches revealed that the two PiSSPs were surprisingly conserved (Figure 5, (Figure S21, Figure S22). Homologs of 1348401 are present throughout the Agaricales and the family strongly expanded in *Armillaria*, with on average 7 genes per species compared to in other Agaricales (Figure S21). The non-pathogenic *A. ectypa* had the fewest genes (3) in this family. We cloned both 1165297 and 1348401 and expressed them transiently in two plant systems: the non-host *Nicotiana benthamiana* and the host plant *E. grandis.* In the former case, expression of 1165297 led to a mild yellowing around the injection site, indicating a hypersensitive response, while 1348401 did not affect the leaf health (Figure 5 B). The expression of these two proteins in *E. grandis* leaves, however, led to rapid cell death within 1 week of agroinfiltration (Figure 5 B), indicating a specific interaction with plant tissue. These data indicate that certain proteins of *A. luteobubalina* can cause necrotic lesions in host plant tissue. The evolutionary conservation of these two proteins outside *Armillaria* is somewhat surprising and is in contrast with observations on other (mostly Ascomycota) effectors ^46, 47^. Whether they trigger cell death via direct (as true effectors) or indirect routes remains to be established. These data raise the possibility that *Armillaria* use specific mechanisms for necrotrophy, as postulated by the gene-for-gene model ^39^.

**Figure 5:**
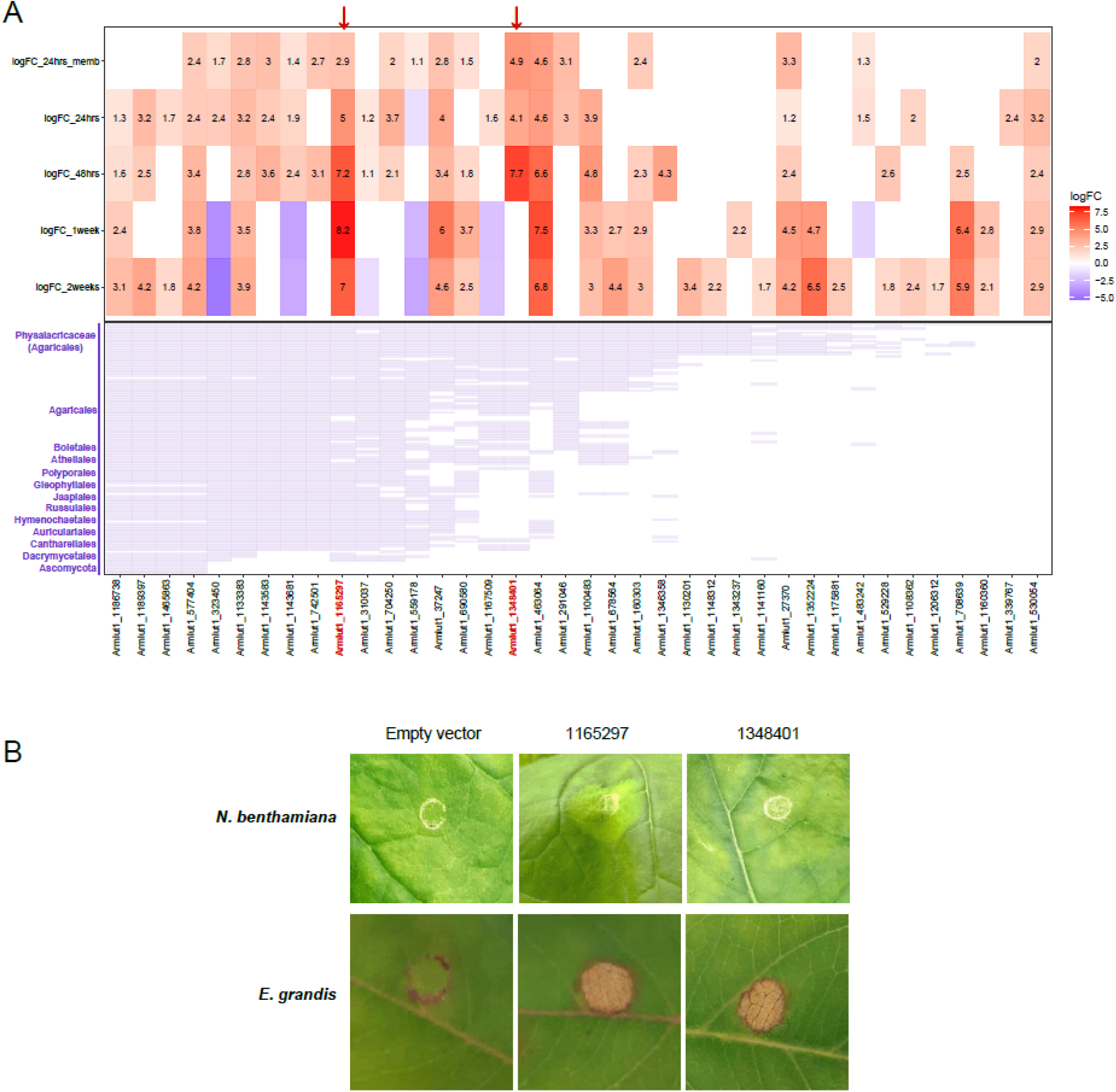
Pathogenicity-induced SSP (PiSSPs) expression and conservation in *A. luteobubalina*. A) Heatmap shows log_2_ fold changes for SSPs, upregulated in at least one time point. Red shows higher and blue depicts lower logFC, followed by presence/absence matrix of homologs of unannotated SSPs in 131 species (Dataset 2). X-axis shows Protein IDs for both heatmap and presence/absence matrix. The two experimentally validated PiSSPs are shown by red arrows. Y-axis of heatmap shows sample comparisons; and species order in presence/absence matrix. B) Transient transformation of the non-host *N. benthamiana* and the host *E. grandis* with either an empty vector, or with an *in planta* expression vector encoding *1165297* or *1348401*.

#### Potential virulence factors in *Armillaria* spp

*In vitro* invasion assays using freshly cut conifer stems were set up to explore gene expression in highly and less-virulent isolates ^23, 24^ of the conifer-specific *A. ostoyae* and *A. borealis* (Figure S12). These experiments emulated cambium colonization and killing, and virulent strains consistently colonized the stems faster. We performed differential gene expression analyses between control and invasive mycelia as well as between virulent and less-virulent strains of the same species (Table S5). SuperSeq ^83^ indicated that the data allowed the detection of up to 89.4-98.6% of all DEGs (Figure S16).

To uncover cambium colonization strategies shared by all strains, we scrutinized genes upregulated in invasive mycelia, relative to control mycelia (Supplementary Note 1). We identified upregulation of PCWDEs, in particular pectinases, as a major transcriptional response to fresh stems in both species. Enrichment ratios confirmed that pectin-related genes are induced in the largest proportions, followed by hemicellulases, cellulases and lignin-related genes (Figure 4). Horizontally transferred intradiol ring-cleavage dioxygenases were also upregulated in these experiments.

We also aimed to identify genes encoding potential virulence factors by comparing highly and less-virulent isolates (Supplementary Note 1). Based on manual curation and sequence-based comparison of upregulated genes against PHI-base ^84^, we identified 303 and 56 candidate virulence factor genes from *A. borealis* and *A. ostoyae* (Table S6). We further determined the secreted proteins to predict potential plant interacting factors. From the *A. borealis* data, 40 secreted proteins, including 9 SSPs, 5 CAZymes, 2 peptidases, 2 intradiol ring-cleavage dioxygenases and one hydrophobin-related gene could be identified. In *A. ostoyae* we found 18 genes, including 3 SSPs, 5 CAZymes and 1-1 genes representing hydrophobins, intradiol ring-cleavage dioxygenases and tyrosinases (Table S6). Notably, highly virulent strains of both species upregulated a pair of polygalacturonase (GH28) and cellulase (GH3) genes, which were both homologous to virulence effectors reported from Ascomycota ^85–87^ (Table S6).

## Discussion

The genus *Armillaria* is a globally important group of primarily tree pathogens that evolved a range of unique traits, from extreme colony sizes to bioluminescence and represent an independent origin of broad host-range necrotrophy. In search of the genetic underpinnings of the unique biology of these fungi, in this study we combined new genomes with transcriptomic profiling of *in planta* and *in vitro* pathosystems. Comparative genomic analyses support the view that *Armillaria* spp. have an expanded protein-coding repertoire and possess a complete set of CAZymes for degrading lignocellulosic plant biomass (including lignin) ^3, 17, 26, 53, 88^. However, we also detected a considerable influx of Ascomycete genes through HGT which, based on the expression data, seem to have influenced both the plant biomass degrading ability and pathogenic attributes of *Armillaria*. Previous studies showed that wood-decay by *Armillaria* is reminiscent of soft-rot ^55, 56, 58, 59^, a lifestyle known only in Ascomycota. Congruently, our phylogenetic PCA analyses separated *Armillaria* from white rot species, which is paradoxical given that white rot is the dominant wood-decay strategy in the Agaricales ^89^. Alien CAZyme genes acquired through HGT from Ascomycota may explain this controversy and establish HGT as a source of novelty in the evolution of fungal plant biomass degrading systems in the Basidiomycota.

*Armillaria* are necrotrophic pathogens that first kill, then feed on their hosts ^1, 17^. Our data provided the first insights into the molecular mechanisms of early- and late stages of colonization of host plants. We detected genes related to four major necrotrophic processes: host immune evasion and suppression, oxidative stress, detoxification, and cytotoxicity, as well as specific regulation of horizontally transferred genes. The presence of pathogenicity-induced small-secreted proteins and the lack of broad plant biomass degrading CAZyme upregulation during early colonization supports the emerging view that, instead of a non-specific, brute-force attack with a broad battery of CAZymes, necrotrophs use specific effectors for modulating the plant immune system ^39, 90^. The expression patterns of the two PiSSPs we described are consistent with the gene-for-gene theory of pathogenicity, which has traditionally been discussed for (hemi-)biotrophs ^91^, but may apply to necrotrophs as well ^39^. Determining whether the two PiSSPs are truly specific effectors will require uncovering the mechanisms of their action, but these data provide an entry point into investigating fine details of *Armillaria* - plant interactions.

In contrast to early-stage infection, cambium colonization was characterized by a broad upregulation of plant cell wall degrading CAZymes. Comparisons of low and highly-virulent strains in these assays further provided insights into genes associated with virulence, such as certain polygalacturonase- or cellulase-encoding genes.

Overall, this study revealed the molecular underpinnings of the lifestyle of a group of widespread pathogens of woody plants, which, combined with existing and emerging experimental tools ^21, 92, 93^ will facilitate research on the molecular mechanisms of plant colonization. We propose that genome evolution in *Armillaria* relied on combined effects of multiple types of genetic innovation, including HGT, and that genes gained early during the evolution of the genus were integrated into cellular regulatory networks of plant biomass degradation and pathogenicity.

## Acknowledgments

We acknowledge support by the ’Momentum’ program of the Hungarian Academy of Sciences (contract no. LP2019-13/2019 to LGN) the European Research Council (grant no. 758161 to LGN) as well as the Eotvos Lorand Research Network (SA-109/2021). GS acknowledges support by the Hungarian National Research, Development, and Innovation Office (GINOP-2.3.2-15-2016-00052). The work (proposals: 10.46936/10.25585/60001060 and 10.46936/10.25585/60001019) conducted by the U.S. Department of Energy Joint Genome Institute (https://ror.org/04xm1d337), a DOE Office of Science User Facility, is supported by the Office of Science of the U.S. Department of Energy operated under Contract No. DE-AC02-05CH11231. Ian Hood and Pam Taylor (Scion Research, New Zealand Forest Research Institute Ltd) kindly provided the *A. nova-zealandiae* 2840 strain. Daniel Lindner (Forest Products Laboratory, USA) kindly shared strains of *A. borealis* and *A. ectypa* for sequencing.

We appreciate the permissions of Gregory Bonito, for using the genome of *Flagelloscypha* sp.

## Methods

### Strains for Sequencing, Assembly and Annotation

*Armillaria* strains (Table S1) were inoculated on Malt Extract Agar (MEA) and incubated at 25°C in the dark for 7-10 days. Identity of each strain was confirmed by amplifying and sequencing the fungal internal transcribed spacer (ITS1-5,8S-ITS2) region and bacterial contamination was checked using universal 16S primers. Strains were inoculated in Malt Extract Broth in 500ml Erlenmeyer flasks and incubated at 25°C in the dark for 4-5 weeks until a substantial quantity of fungal biomass was grown. Fungal mass was stored at -80°C until DNA and RNA extraction. Prior to nucleic acid extraction, fungal tissues were homogenized using liquid nitrogen in mortar and pestle. DNA extraction was performed using the Blood & Cell Culture DNA Maxi Kit (Qiagen Inc.) and RNA extraction was performed using the RNeasy Midi Kit (Qiagen Inc.) as per manufacturer’s instructions. Using the DNA we again confirmed strain identity by sequencing the ITS region and nucleic acid quantity was measured using Qubit (ThermoFisher) according to the manufacturer’s instructions.

### Genomic and transcriptomic library preparation, sequencing, assembly and annotation

For *A. nabsnona*, *A. mellea*, and *A. ectypa*, 5 ug of genomic DNA was sheared to approximately 15-20 kb using Megaruptor3 (Diagenode). The sheared DNA was treated with DNA Prep to remove single-stranded ends, and DNA damage repair mix followed by end repair, A-tail and ligation of PacBio overhang adapters using SMRTbell Express Template Prep 2.0 Kit (Pacific Biosciences). The final libraries were size selected with BluePippin (Sage Science) at 10 kb cutoff size. For *A. borealis, A. fumosa, A. novae-zelandiae*, and *A. tabescens*, 5 ug of genomic DNA was left unsheared due to marginal HMW DNA quality. The unsheared gDNA was treated with exonuclease to remove single-stranded ends and DNA damage repair mix followed by end repair and ligation of blunt adapters using SMRTbell Template Prep Kit 1.0 (Pacific Biosciences). All libraries were purified with AMPure PB beads.

The PacBio Sequencing primers were then annealed to the SMRTbell template libraries and sequencing polymerase was bound to them using Sequel Binding kit 2.0 (*A. nabsnona*, *A. mellea*, *A. tabescens*, and *A. ectypa*), 2.1 (*A. borealis*), or 3.0 (*A. fumosa* and *A. novae-zelandiae*). The prepared SMRTbell template libraries were then sequenced on a Pacific Biosciences’ Sequel sequencer using v3 sequencing primer, 1M v2 (*A. borealis*, *A. nabsnona*, *A. mellea*, *A. tabescens*, and *A. ectypa*) or v3 (*A. fumosa*, and *A. novae-zelandiae*) SMRT cells, and Version 2.1 (*A. borealis*, *A. nabsnona*, *A. mellea*, *A. tabescens*, and *A. ectypa*) or 3.0 (*A. fumosa* and *A. novae-zelandiae*) sequencing chemistry with 1×360 & 1×600 sequencing movie run times.

Filtered subread data was processed to remove artifacts and assembled together with Falcon (https://github.com/PacificBiosciences/FALCON) version 1.8.8 (*A. ectypa*, *A. nabsnona*, *A. tabescens*, and *A. borealis*) or version pb-assembly 0.0.2, falcon-kit 1.2.3, pypeflow 2.1.0 (*A. mellea*, *A. novae-zelandiae*, and *A. fumosa*) to generate an initial assembly. Mitochondria was assembled separately from the Falcon pre-assembled reads (preads) using an in-house tool (assemblemito.sh (*A. ectypa*, *A. nabsnona*, *A. tabescens*, *A. borealis*, and *A. mellea*) or assemblemito.py (*A. fumosa* and *A. novae-zelandiae*)), used to filter the preads, and polished with Arrow version SMRTLink v5.0.1.9578 (*A. ectypa* and *A. nabsnona*), v5.1.0.26412 (*A. tabescens* and *A. borealis*), v6.0.0.47841 (*A. mellea*) or v7.0.1.66975 (*A. novae-zelandiae* and *A. fumosa*) (https://github.com/PacificBiosciences/GenomicConsensus). A secondary Falcon assembly was generated using the mitochondria-filtered preads, improved with finisherSC ^94^ version 2.0 (*A. ectypa* and *A. nabsnona*) or 2.1 (*A. fumosa*), and polished with Arrow version SMRTLink v5.0.1.9578 (*A. ectypa* and *A. nabsnona*), v5.1.0.26412 (*A. tabescens* and *A. borealis*), v6.0.0.47841 (*A. mellea*), or v7.0.1.66975 (*A. novae-zelandiae* and *A. fumosa*). Contigs less than 1000 bp were excluded.

Stranded cDNA libraries were generated using the Illumina Truseq Stranded mRNA Library Prep kit. mRNA was purified from 200 ng (*A. nabsnona*) or 1ug (all other species) of total RNA using magnetic beads containing poly-T oligos. For *A. nabsnona*, mRNA was fragmented using divalent cations and high temperature. The fragmented RNA was reverse transcribed using random hexamers and SSII (Invitrogen) followed by second strand synthesis. For all other species, mRNA was fragmented and reverse transcribed using random hexamers and SSII (Invitrogen) followed by second strand synthesis. The fragmented cDNA was treated with end-pair, A-tailing, adapter ligation, and 10 (*A. nabsnona*) or 8 (all other species) cycles of PCR.

All prepared libraries were quantified using KAPA Biosystems’ next-generation sequencing library qPCR kit and run on a Roche LightCycler 480 real-time PCR instrument. Sequencing was performed on the Illumina NovaSeq sequencer using NovaSeq XP V1 reagent kits, S4 flowcell (*A. fumosa*, *A. novae-zelandiae*, and *A. mellea*) or on the Illumina HiSeq2500 sequencer using TruSeq paired-end cluster kits v4 and TruSeq SBS sequencing kits v4 (*A. nabsnona*, *A. borealis*, *A. tabescens* and *A. ectypa*), following a 2×150 indexed run recipe.

Raw reads were evaluated with BBDuk (https://sourceforge.net/projects/bbmap/) for artifact sequence by kmer matching (kmer=25), allowing 1 mismatch and detected artifacts were trimmed from the 3’ end of the reads. RNA spike-in reads, PhiX reads, and reads containing any Ns were removed. Quality trimming was performed using the phred trimming method set at Q6. Following trimming, reads under the length threshold were removed (minimum length 25 bases or 1/3 of the original read length - whichever is longer). Filtered reads were assembled into consensus sequences using Trinity (ver. 2.3.2) ^95^ with the --normalize_reads and --jaccard_clip options.

Genomes were annotated with the support of their corresponding transcriptomes using the JGI Annotation Pipeline. Assemblies and annotations are available from the JGI Genome Portal MycoCosm (https://mycocosm.jgi.doe.gov) ^96, 97^.

### Taxon sampling and Functional Annotations

We used 3 datasets in this study. Dataset1 comprises 66 species (64 Agaricales, 2 Boletales outgroups), which were used to analyze gene family evolution in *Armillaria* (Table S1: Dataset1). This was extended into a phylogenetically more diverse Dataset2, which was used to perform gene copy number analysis of wood-decay gene repertoires and comprised 131 fungal species from both Basidiomycota and Ascomycota, including species with various lifestyles such as white-rotters, brown-rotters, litter-decomposers, ECM fungi, pathogens, and species with uncertain ecologies (Table S1: Dataset2). Dataset3 was used to identify horizontally transferred genes and comprised 942 species, including species ranging from fungi, bacteria as well as plants (Table S1: Dataset3). All the species used in these datasets are published and publicly available sequences from JGI Mycocosm (except a few that were used with Author’s permission). Proteomes of all the species used in this manuscript were functionally annotated by using InterPro (IPR) Scan (v.5.46-81.0) ^98^. Secreted and small secreted (at least 300 amino acids) proteins were identified as described previously ^99^.The luciferase cluster and its synteny were identified in *Armillaria* species and 5 Physalacriaceae outgroups as described previously^49^. Orthologous groups from these genomes were inferred using OrthoFinder (version 2.5.4) ^100, 101^ and synteny around the luciferase cluster was visualized based on ordering and orienting scaffolds using the *Armillaria ostoyae* C18/9 genome and using genoPlotR ^102^ package. The hispidin synthase and cytochrome P450 gene were previously missing and re-annotated on the scaffold of NODE_104435 in *Armillaria mellea* DSM 3731 using predictions of *Armillaria mellea* ELDO17 v1.0.

### Genome evolution in *Armillaria* spp

Reconstruction of genome wide gene family evolution was performed using Dataset1. Predicted protein sequences were clustered into orthogroups using OrthoFinder (v2.3.1) ^100, 101^ with default settings. For the species tree, multiple sequence alignment for 81 single copy orthogroups (SCOGs) were inferred using MAFFT (-auto) ^103^, followed by removal of gapped regions from the alignments using TrimAl ^104^ with a gap threshold of 0.9. Trimmed alignments were then concatenated into a supermatix using an in-house R-script. The species tree was reconstructed in RAxML (v8.2.12) ^105^ with the option of rapid bootstrap analysis and search for the best-scoring ML tree under the PROTGAMMAWAG model of protein evolution. The tree was rooted using FigTree and is provided in Table S1.

We analyzed the genome-wide duplications and losses across gene families using the COMPARE method ^106^ which uses reconciled gene trees to perform Dollo parsimony mapping and ortholog coding to tell apart duplications from speciation events. To obtain the reconciled gene trees for COMPARE, we first inferred multiple sequence alignments for the Orthofinder orthogroups with at least 4 proteins using MAFFT (-auto) ^103^. Aligned clusters were trimmed to remove spurious regions using TrimAL ^104^ with -strict parameter. Clusters for which TrimAL resulted in blank alignments (due to -strict parameter) were used in their non-trimmed form. In total, 16,936 trimmed and 819 non-trimmed clusters were used in RAxML (v8.2.12) ^105^ for gene tree inference, followed by Shimodaira-Hasegawa (sh)-like branch support calculation also in RAxML under the PROTGAMMAWAG substitution model. The 17,755 gene trees with their SH-like support values, along with the species tree were used for rerooting, followed by gene tree species tree reconciliation in Notung (v2.9) run in batch mode with edge weight threshold set to 70. Reconciled gene trees were used to reconstruct the gene duplication/loss scenarios in orthogroups along the species tree using the COMPARE pipeline (available at https://github.com/laszlognagy/COMPARE). For each gene family, the number of gains, losses, net gains (sum of gains and losses), and ancestral copy numbers were obtained and their summary was mapped back onto the species tree. GO terms significantly enriched (p-values < 0.05) among the duplications at *Armillaria* MRCA were identified by topGO ^42^ using the weight01 algorithm and Fisher testing.

### Analyses of CAZymes in *Armillaria* spp

Dataset2 was used to analyze where Physalacriaceae (including *Armillaria*) fits in terms of their wood-decay repertoires. Annotation of Carbohydrate Active enzymes (CAZymes) for the required species were downloaded from JGI Mycocosm (species whose CAZymes were not present in JGI Mycocosm were annotated using the CAZy annotation pipeline ^107^. Based on previously published reports, annotated CAZymes were further categorized based on their putative plant cell wall degrading preferences (Table S3) into those acting on cellulose, hemicellulose, pectin and lignin. We subjected the proteomes of these 131 species to OrthoFinder (v2.3.1) ^100, 101^ with default settings, giving us a total of 41,205 orthogroups. For inferring the species tree, we generated SCOGs using a set of in-house R scripts which gave us 548 orthogroups in which the duplications were caused by gains at species levels (or terminal gains). From these we omitted proteins showing less similarity based on amino acid distance against the other members of the cluster. In this step we used at least 40% of all species for each cluster. Finally, the resulting 514 SCOGs were concatenated (min 60 length of AA, and at least 66 species) into one supermatrix and followed by species tree inference in IQ-Tree ^108^.

We generated phylogenetic PCA using the phyl.pca ^109^ function from Phytools ^110^ for Physalacriaceae along with WR and LD fungi. We compare *Armillaria* and other Physalacriaceae to only WR and LD fungi, because including brown rot and mycorrhizal species, which have huge differences in gene content, reduced the resolution of the plots. We provided the function with a gene copy number matrix normalized according to proteome sizes for each substrate category and the above described ML species tree (subsetted to WR, LD and Physalacriaceae species) as inputs. Independent contrasts were calculated under the Brownian motion model and the parameter mode=“cov”. The substrate-wise gene copy numbers and their resulting phylogenetic PCA loadings are provided in Table S3. We also plotted their proteome-size normalized copy numbers into barplots for visualization of copy numbers across the phylogeny (Figure S9, Figure S6, Figure S10, Figure S7, Figure S8).

We further investigated if Physalacriaceae members have CAZymes shared with Ascomycota. For checking this hypothesis, we first identified the CAZy orthogroups from the 131 species, by selecting orthogroups with at least 50% of the proteins annotated as CAZymes, and at least 5 proteins. This gave us 401 CAZy orthogroups which were used to perform Fisher’s exact test-based enrichment for identifying co-enriched CAZy families in Physalacriaceae and Ascomycota with respect to WR and LD (Table S3).

### Analyses of horizontally transferred genes

In order to check if Physalacriaceae species are more similar to Ascomycota than expected by chance, which could either come from horizontal gene transfer or long retained ancestral genes, we compiled the Dataset3. This comprised a taxonomically diverse dataset with a total of 942 species, ranging from 110 fungi from Basidiomycota (including 20 Physalacriaceae), 741 from Ascomycota, 26 from Mucoromycota, 10 from Zoopagomycota, 13 from Chytridiomycota and 17 other early-diverging fungi as well as 15 bacterial and 10 plant species.

We identified candidate horizontally transferred (HT) genes using the Alien index (AI), for which we ran MMseqs (easy-search, e-value 0.001) ^111^ using all Physalacriaceae proteomes merged together as our query, against a database of proteomes of all the other species as our subject. Using an in-house R script, we parsed the output to retain only the top hits (based on e-values) for each taxonomic group. AI scores were calculated as [log_10_(best hit to species within the group lineage + 1×10^-200^) - log_10_(best hit to species outside the group lineage + 1×10^-200^)]112,113.

Further, for phylogenetic validation of HT events, we used a subset of 329 species from Dataset3 based on AI of top HT donors. In this case, we restricted our donor list to major HT contributors only, *viz.* Ascomycota, followed by Mucoromycota and Zoopagomycota as outgroups. The proteomes of these 329 species were subjected to OrthoFinder (v2.5.4) ^100, 101^ clustering with default settings. From these, we fished out the orthogroups with at least one AI-based HT gene and also the orthogroups containing CAZymes. Further, we retained orthogroups having at least one Physalacriaceae and one Ascomycota/Mucoromycota/Zoopagomycotina species, resulting in a total of 675 orthogroups.

Proteins from these 675 orthogroups were aligned using MAFFT (-auto) ^103^ TrimAL (-strict)^104^. Trimmed alignments were used to infer gene trees using IQ-Tree^108^ with ultrafast bootstrap (1000 replicates) under the WAG+G substitution model. Using an in-house R-script, we identified potential HT events by extracting clades from the gene trees based on support values (>70%) and taxon occupancy (receiver clade with >70% Physalacriacae species, donor clade with >70% Ascomycota, and finally sister clade with >70% Ascomycota to ensure the direction of gene transfer). Physalacriceae proteins from these putative HT clades, were used as a query against the UniRef100 database ^61^ from the UniProt Reference Cluster using MMSeqs easy-search, e-value 0.001) ^111^. To remove the low sequence similarity matches and false hits, we retained hits with percent identity and bidirectional coverage of more than 45%. We parsed the filtered hits to retain only the top 100 hits (based on percent identity) for each query, and the taxonomic category contributing predominantly among the top 100 hits was assigned as the putative donor. Further, we classified the assigned donor into two categories: Type A, where the top 100 hits were dominated and confidently associated with Ascomycota, and Type B, where the top 100 hits were dominated by non-Agaricomycetes species, however ambiguously distributed among different taxa (Table S4).

### Live plant and stem invasion pathosystems

To understand gene expression in *Armillaria* and its host during the early stages of host infection, we performed live *in planta* assays between *E. grandis* and *A. luteobubalina. A. luteobubalina* was cultured onto half-strength modified Melin–Norkrans (MMN) media (pH 5.5; 1g l^−1^ glucose) and grown for one month in the dark. *E. grandis* seeds were sterilized in 30% hydrogen peroxide (H_2_O_2_, v/v) and germinated on 1% (w/v) water agar one month (25°C; 16 h light cycle). Four weeks before contact with the fungus, seedlings were transferred onto half-strength MMN medium ((pH 5.5; 1 g l^−1^ glucose) and grown at 22–30°C night/day temperature with a 16 h light cycle. Once the *E. grandis* seedlings were two months old and the fungal cultures had grown for one month, the plants were separated into one of three treatment categories: (1) fungal-free controls whereby the seedlings were transferred onto new half-strength MMN medium; (2) ‘pre-symbiosis’ which involved the transfer of seedlings onto new half-strength MMN medium in indirect contact with fungal mycelium for 24 h by separating the two organisms by a permeable cellophane membrane; (3) ‘physical contact’ seedlings were transferred onto new half-strength MMN medium and then placed into direct contact with the active growing edge of a fungal colony and then samples were harvested at 24h, 48 h, 1 week, and 2 week post-contact. These plates were then closed using micropore tape to allow for gas exchange with the external environment. Four biological replicates per treatment and timepoint were generated and harvested at the described timepoint into liquid nitrogen and stored at -80°C until RNA extraction.

For stem-invasion assays, high and low virulent isolates of conifer-specific *A. ostoyae* (C18 - highly virulent, C2 - low virulent), and *A. borealis* (A6 - highly virulent, A4 - low virulent) ^23, 114^ were maintained in the dark on Petri dishes on RS medium (40g malt extract, 20g dextrose, 5g bacto peptone, 19g agar / 1L) at 24°C and 4°C, respectively. Before the start of the experiment, fresh cultures of all isolates were set up. The system for growing subcortical mycelial fans in the laboratory consisted of plastic jars containing a layer of moistened and inoculated RSTO medium ^17^. After approximately 10 days of incubation in dark at 24°C, 10cm long, freshly cut stem segments of Norway spruce were placed on top of this growing mycelial lawn, allowing *Armillaria* to invade the stem segments. The timing and effectiveness of the advancing invasive mycelial fans were followed by their arrival in small monitoring “windows” (1 x 1.5 cm) cut into the bark halfway up to the top of the stems. The samples were harvested soon after the mycelial front line appeared in the cutouts. *Armillaria* vegetative mycelia collected from the jars without spruce stems were used as controls.

### RNA isolation and sequencing

For *in planta* assays, live tissues, frozen samples were harvested and extracted using the ISOLATE II Plant miRNA kit (Bioline, Sydney, Australia) as per the manufacturer’s instructions. Following extraction, the RNA samples were sequenced at the Beijing Genome Institute (BGI). For the stem invasion assays, sections of the mycelial fan that were collected from under the bark of infected colonized stems were frozen in liquid nitrogen and stored at -80°C. RNA was isolated from the frozen samples using the RNeasy Plant Midi Kit (Qiagen Inc.) according to the manufacturer’s protocol. Prior to the RNA isolation, fungal tissues were homogenized with the help of liquid nitrogen, mortar and pestle. RNA quantity was measured using Qubit (ThermoFisher) according to the manufacturer’s protocol. Biological triplicates were analyzed for all sample types. The libraries for Illumina sequencing were prepared using NEBNext Ultra II Directional RNA Library Prep Kit for Illumina (NEB, Ipswitch, MA, USA). Briefly, 100 ng RNA was enriched using RiboCop rRNA Depletion Kits (Lexogen, Austria). Thereafter, the RNA was fragmented, end prepped and adapter-ligated. Finally, the libraries were amplified according to the manufacturer’s instructions. The quality of the libraries was checked on Agilent 4200 TapeSation System using D1000 Screen Tape (Agilent Technologies, Palo Alto, CA, USA), the quantity was measured on Qubit 3.0. Illumina sequencing was performed on the NovaSeq 6000 instrument (Illumina, San Diego, CA, USA) with 2 × 151 run configuration. Raw RNA-Seq reads were aligned against the *A. ostoyae* (NCBI genome GCA_900157425.1 version 2) and *A. borealis* (JGI: *Armillaria borealis* FPL87.14 v1.0) genomes using STAR v2.7 .5a ^115^. After alignment, the level of expression was estimated using RSEM v1.3.1 ^116^.

### Analysis of expression data

For the stem invasion assay, the estimated counts matrix was used for differential expression analysis using Limma-Voom ^117^. Prior to running the differential expression analysis, counts were normalized using TMM (trimmed mean of M values) using edgeR v3.38.1 ^118^ and lowly expressed genes (cpm ≤ 10) were filtered. Differential gene expression analysis for the *in-planta* pathosystem data were carried out as our previous study ^58^.

In both analyses, genes with at least two-fold expression change and FDR value <=0.05 were considered significant. Multidimensional scaling (“plotMDS” function in edgeR) was applied to visualize gene expression profiles. Expression heatmaps were generated by hierarchically clustering based on average linkage of FPKM values with heatmap.2 function from gplots package

To check reliability of both expression data, we used the superSeq package ^83^ which uses a subsampling approach to simulate and predict the number of differentially expressed genes at lower read depths and random sampling points from the original dataset. These subsampled reads are then extrapolated to predict the relationship between statistical power and read depth.

Genes from the time-series *in planta* assay from both fungal and the plant side were clustered into co-expression based modules using STEM (v1.3.11) ^64^. GO terms significantly enriched (p-values < 0.05) among each of the STEM modules were identified by topGO ^42^, using the weight01 algorithm and Fisher testing.

### Experimental validation of Pathogenicity-induced SSPs

Aliquots of RNA extracted from the *in planta* pathosystem assay (6 timepoints, 24 samples) were pooled and used for generation of full length cDNA using the Tetro cDNA synthesis kit (Bioline) according to manufacturer’s instructions and using only the oligo-dT primer. Two *A. luteobubalina* HWK02 v1.0 SSP sequences were amplified using the KAPA HiFi Hotstart Readymix (Roche) according the manufacturer’s instructions using the following primers: *Armlut1_1165297*(5’-GGGGACAAGTTTGTACAAAAAAGCAGGCTTAATGTTYCARYTNYTNTTYGCNGC-3’ and 5’-GGGGACCACTTTGTACAAGAAAGCTGGGTGTTACTCTTCTAGCAGCTTCACGTC-3’) and *Armlut_1348401*(5’-GGGGACAAGTTTGTACAAAAAAGCAGGCTTAATGTTGTTCAGTTTCTTCCTCTTCTACC-3’ and 5’-GGGGACCACTTTGTACAAGAAAGCTGGGTGCTARTCNACRCANGTNGTNSWDATRTCNG-3’). Successful amplification was verified using gel electrophoresis, products were gel purified using the Wizard SV Gel and PCR Cleanup kit (Promega) and ligated into pDONR221-b1b4 (Invitrogen) using BP ligase (Invitrogen). Ligations were transformed into *E. coli* and positive colonies were selected based on PCR screening. Positive bacterial colonies were grown overnight in LB + kanamycin (50 mg/L) and plasmids purified using the PureLink HiPure Plasmid Miniprep kit. Gene inserts were sequence verified using Sanger sequencing at the Hawkesbury Institute for the Environment sequencing platform. Plasmids containing the proper gene insert were used in an LR ligation with pBiFCt-2in1destination vector with an empty cassette ligated into the b2b3 position. Similarly, as a control for *in planta* transformations, a version of pBIFCt-2in1 with empty cassettes was generated as an empty vector control.

Screening procedures following were as per with pDONR221. Positive plasmids were purified and transformed into *Agrobacterium tumefaciens* GV3101. Expression in *N. benthamiana* was performed as per Plett et al., 2020 . Similarly, young leaves of 6 month old *E. grandis* saplings grown at 25°C constant temperature (16hr light/8hr dark) were agroinfiltrated using the same solutions as for *N. benthamiana.* As opposed to *N. benthamiana* leaves, *E. grandis* leaves were more recalcitrant to infiltration without damaging the leaves, therefore the bacteria was only introduced into a circle the size of the syringe bore. Both plant systems were left for seven days and the leaves photographed at the site of infiltration. Leaves from three separate plants were infiltrated for each plant model system and representative images presented.

## Notes

### Competing Interest Statement

The authors have declared no competing interest.

### Summary of Updates

The author's name Boris Indjic was corrected to Boris Indic in the metadata. All other files remain the same.

## References

1. Baumgartner, K., Coetzee, M. P. A. & Hoffmeister, D. Secrets of the subterranean pathosystem of *Armillaria*: Subterranean pathosystem of *Armillaria*. Mol. Plant Pathol. 12, 515–534 (2011).

2. Heinzelmann, R. et al. Latest advances and future perspectives in *Armillaria* research. Can. J. Plant Pathol. 41, 1–23 (2019).

3. Sipos, G., Anderson, J. B. & Nagy, L. G. Armillaria. Curr. Biol. 28, PR297-R298 (2018).

4. Coetzee, M., Wingfield, B. & Wingfield, M. Armillaria Root-Rot Pathogens: Species Boundaries and Global Distribution. Pathogens 7, 83 (2018).

5. Shaw, C. G. & Roth, L. F. Control of Armillaria root rot in managed coniferous forests.: A literature review. For. Pathol. 8, 163–174 (1978).

6. Drakulic, J., Gorton, C., Perez-Sierra, A., Clover, G. & Beal, L. Associations Between *Armillaria* Species and Host Plants in U.K. Gardens. Plant Dis. 101, 1903–1909 (2017).

7. Raabe, R. D. Host list of the root rot fungus, *Armillaria mellea*. Hilgardia 33, 25–88 (1962).

8. Baumgartner, K. Root Collar Excavation for Postinfection Control of Armillaria Root Disease of Grapevine. Plant Dis. 88, 1235–1240 (2004).

9. Cleary, M., Morrison, D. J. & Kamp, B. Symptom development and mortality rates caused by *Armillaria ostoyae* in juvenile mixed conifer stands in British Columbia’s southern interior region. For. Pathol. 51, (2021).

10. Murray, M. P. & Leslie, A. Climate, radial growth, and mortality associated with conifer regeneration infected by root disease (*Armillaria ostoyae)*. For. Chron. 97, 43–51 (2021).

11. Anderson, J. B. & Catona, S. Genomewide mutation dynamic within a long-lived individual of *Armillaria gallica*. Mycologia 106, 642–648 (2014).

12. Anderson, J. B. et al. Clonal evolution and genome stability in a 2500-year-old fungal individual. Proc. R. Soc. B Biol. Sci. 285, 20182233 (2018).

13. Smith, M. L., Bruhn, J. N. & Anderson, J. B. The fungus Armillaria bulbosa is among the largest and oldest living organisms. Nature 356, 428–431 (1992).

14. Ullrich, R. C. & Anderson, J. B. Sex and diploidy in Armillaria mellea. Exp. Mycol. 2, 119–129 (1978).

15. Mihail, J. D. & Bruhn, J. N. Dynamics of bioluminescence by *Armillaria gallica*, A. mellea and A. tabescens. Mycologia 99, 341–350 (2007).

16. Porter, D. L. et al. The melanized layer of Armillaria ostoyae rhizomorphs: Its protective role and functions. J. Mech. Behav. Biomed. Mater. 125, 104934 (2022).

17. Sipos, G. et al. Genome expansion and lineage-specific genetic innovations in the forest pathogenic fungi Armillaria. Nat. Ecol. Evol. 1, 1931–1941 (2017).

18. Koch, R. A. et al. Symbiotic nitrogen fixation in the reproductive structures of a basidiomycete fungus. Curr. Biol. 31, 3905–3914.e6 (2021).

19. Devkota, P. & Hammerschmidt, R. The infection process of Armillaria mellea and Armillaria solidipes. Physiol. Mol. Plant Pathol. 112, 101543 (2020).

20. Wong, J. W. H. et al. Comparative metabolomics implicates threitol as a fungal signal supporting colonization of *Armillaria luteobubalina* on eucalypt roots. Plant Cell Environ. 43, 374–386 (2020).

21. Ford, K. L., Henricot, B., Baumgartner, K., Bailey, A. M. & Foster, G. D. A faster inoculation assay for *Armillaria* using herbaceous plants. J. Hortic. Sci. Biotechnol. 92, 39– 47 (2017).

22. Bendel, M., Kienast, F. & Rigling, D. Genetic population structure of three Armillaria species at the landscape scale: a case study from Swiss Pinus mugo forests. Mycol. Res. 110, 705–712 (2006).

23. Heinzelmann, R., Prospero, S. & Rigling, D. Virulence and Stump Colonization Ability of *Armillaria borealis* on Norway Spruce Seedlings in Comparison to Sympatric *Armillaria* Species. Plant Dis. 101, 470–479 (2017).

24. Prospero, S., Rigling, D. & Holdenrieder, O. Population structure of Armillaria species in managed Norway spruce stands in the Alps. New Phytol. 158, 365–373 (2003).

25. Caballero, J. R. I. et al. Genomic Comparisons of Two Armillaria Species with Different Ecological Behaviors and Their Associated Soil Microbial Communities. Microb. Ecol. (2022) doi:10.1007/s00248-022-01989-8.

26. Collins, C. et al. Genomic and Proteomic Dissection of the Ubiquitous Plant : Toward a New Infection Model System. J. Proteome Res. L 12, 2552–2570 (2013).

27. Heinzelmann, R., Rigling, D., Sipos, G., Münsterkötter, M. & Croll, D. Chromosomal assembly and analyses of genome-wide recombination rates in the forest pathogenic fungus Armillaria ostoyae. Heredity 124, 699–713 (2020).

28. Kedves, O., et al. Epidemiology, Biotic Interactions and Biological Control of Armillarioids in the Northern Hemisphere. Pathogens 10, 76 (2021).

29. Kolesnikova, A. I. et al. Mobile genetic elements explain size variation in the mitochondrial genomes of four closely-related Armillaria species. BMC Genomics 20, 351 (2019).

30. Wingfield, B. D. et al. IMA Genome-F 6: Draft genome sequences of Armillaria fuscipes, Ceratocystiopsis minuta, Ceratocystis adiposa, Endoconidiophora laricicola, E. polonica and Penicillium freii DAOMC 242723. IMA Fungus 7, 217–227 (2016).

31. Kabbage, M., Yarden, O. & Dickman, M. B. Pathogenic attributes of Sclerotinia sclerotiorum : Switching from a biotrophic to necrotrophic lifestyle. Plant Sci. 233, 53–60 (2015).

32. Liang, X. & Rollins, J. A. Mechanisms of Broad Host Range Necrotrophic Pathogenesis in *Sclerotinia sclerotiorum*. Phytopathology® 108, 1128–1140 (2018).

33. Xu, L., Li, G., Jiang, D. & Chen, W. *Sclerotinia sclerotiorum* : An Evaluation of Virulence Theories. Annu. Rev. Phytopathol. 56, 311–338 (2018).

34. O’Connell, R. J. et al. Lifestyle transitions in plant pathogenic *Colletotrichum* fungi deciphered by genome and transcriptome analyses. Nat. Genet. 44, 1060–1065 (2012).

35. Foley, R. C., Kidd, B. N., Hane, J. K., Anderson, J. P. & Singh, K. B. Reactive Oxygen Species Play a Role in the Infection of the Necrotrophic Fungi, Rhizoctonia solani in Wheat. PLOS ONE 11, e0152548 (2016).

36. Newman, T. E. & Derbyshire, M. C. The Evolutionary and Molecular Features of Broad Host-Range Necrotrophy in Plant Pathogenic Fungi. Front. Plant Sci. 11, 591733 (2020).

37. Westrick, N. M., Smith, D. L. & Kabbage, M. Disarming the Host: Detoxification of Plant Defense Compounds During Fungal Necrotrophy. Front. Plant Sci. 12, 651716 (2021).

38. Olson, Å. et al. Insight into trade-off between wood decay and parasitism from the genome of a fungal forest pathogen. New Phytol. 194, 1001–1013 (2012).

39. Shao, D., Smith, D. L., Kabbage, M. & Roth, M. G. Effectors of Plant Necrotrophic Fungi. Front. Plant Sci. 12, 687713 (2021).

40. Chin, C.-S. et al. Phased diploid genome assembly with single-molecule real-time sequencing. Nat. Methods 13, 1050–1054 (2016).

41. Koch, R. A., Wilson, A. W., Séné, O., Henkel, T. W. & Aime, M. C. Resolved phylogeny and biogeography of the root pathogen *Armillaria* and its gasteroid relative, *Guyanagaster*. BMC Evol. Biol. 17, (2017).

42. Alexa, A. & Rahenfuhrer, J. topGO: Enrichement Analysis for Gene Ontology. (2016).

43. Baccelli, I. Cerato-platanin family proteins: one function for multiple biological roles? Front. Plant Sci. 5, (2015).

44. Li, S. et al. The Novel Cerato-Platanin-Like Protein FocCP1 from Fusarium oxysporum Triggers an Immune Response in Plants. Int. J. Mol. Sci. 20, 2849 (2019).

45. Muraosa, Y., Toyotome, T., Yahiro, M. & Kamei, K. Characterisation of novel-cell-wall LysM-domain proteins LdpA and LdpB from the human pathogenic fungus Aspergillus fumigatus. Sci. Rep. 9, 3345 (2019).

46. Plett, J. M. & Plett, K. L. Leveraging genomics to understand the broader role of fungal small secreted proteins in niche colonization and nutrition. ISME Commun. 2, 49 (2022).

47. Lo Presti, L., et al. Fungal Effectors and Plant Susceptibility. Annu. Rev. Plant Biol. 66, 513–545 (2015).

48. Sonah, H., Deshmukh, R. K. & Bélanger, R. R. Computational Prediction of Effector Proteins in Fungi: Opportunities and Challenges. Front. Plant Sci. 7, (2016).

49. Ke, H.-M. et al. *Mycena* genomes resolve the evolution of fungal bioluminescence. Proc. Natl. Acad. Sci. 117, 31267–31277 (2020).

50. Nagy, L. G. et al. Genetic Bases of Fungal White Rot Wood Decay Predicted by Phylogenomic Analysis of Correlated Gene-Phenotype Evolution. Mol. Biol. Evol. 34, 35– 44 (2017).

51. Sun, P. et al. Fungal glycoside hydrolase family 44 xyloglucanases are restricted to the phylum Basidiomycota and show a distinct xyloglucan cleavage pattern. iScience 25, 103666 (2022).

52. Resl, P. et al. Large differences in carbohydrate degradation and transport potential among lichen fungal symbionts. Nat. Commun. 13, 2634 (2022).

53. Collins, C. et al. Proteomic Characterization of *Armillaria mellea* Reveals Oxidative Stress Response Mechanisms and Altered Secondary Metabolism Profiles. Microorganisms 5, 60 (2017).

54. Floudas, D. et al. Evolution of novel wood decay mechanisms in Agaricales revealed by the genome sequences of *Fistulina hepatica* and *Cylindrobasidium torrendii*. Fungal Genet. Biol. 76, 78–92 (2015).

55. Campbell, W. G. The chemistry of the white rots of wood. Biochem. J. 25, 2023– 2027 (1931).

56. Campbell, W. G. The chemistry of the white rots of wood. Biochem. J. 26, 1829– 1838 (1932).

57. Daniel, G., Volc, J. & Nilsson, T. Soft rot and multiple T-branching by the basidiomycete *Oudemansiella mucida*. Mycol. Res. 96, 49–54 (1992).

58. Sahu, N. et al. Hallmarks of Basidiomycete Soft- and White-Rot in Wood-Decay - Omics Data of Two Armillaria Species. Microorganisms 9, 149 (2021).

59. Schwarze, F. W. M. R. Wood decay under the microscope. Fungal Biol. Rev. 21, 133–170 (2007).

60. Gladyshev, E. A., Meselson, M. & Arkhipova, I. R. Massive Horizontal Gene Transfer in Bdelloid Rotifers. Science 320, 1210–1213 (2008).

61. Suzek, B. E. et al. UniRef clusters: a comprehensive and scalable alternative for improving sequence similarity searches. Bioinformatics 31, 926–932 (2015).

62. Worrall, J. J., Anagnost, S. E. & Zabel, R. A. Comparison of Wood Decay among Diverse Lignicolous Fungi. Mycologia 89, 199 (1997).

63. Yang, G. et al. A cerato-platanin protein SsCP1 targets plant PR1 and contributes to virulence of *Sclerotinia sclerotiorum*. New Phytol. 217, 739–755 (2018).

64. Ernst, J. & Bar-Joseph, Z. STEM: a tool for the analysis of short time series gene expression data. BMC Bioinformatics 7, 191 (2006).

65. Bautista, D. et al. Comprehensive Time-Series Analysis of the Gene Expression Profile in a Susceptible Cultivar of Tree Tomato (Solanum betaceum) During the Infection of Phytophthora betacei. Front. Plant Sci. 12, 730251 (2021).

66. Biniaz, Y. et al. Transcriptome Meta-Analysis Identifies Candidate Hub Genes and Pathways of Pathogen Stress Responses in Arabidopsis thaliana. Biology 11, 1155 (2022).

67. Ge, Y. et al. Transcriptome analysis identifies differentially expressed genes in maize leaf tissues in response to elevated atmospheric [CO _2_]. J. Plant Interact. 13, 373– 379 (2018).

68. Chen, H., Quintana, J., Kovalchuk, A., Ubhayasekera, W. & Asiegbu, F. O. A cerato-platanin-like protein HaCPL2 from Heterobasidion annosum sensu stricto induces cell death in Nicotiana tabacum and Pinus sylvestris. Fungal Genet. Biol. 84, 41–51 (2015).

69. Curir, P., Dolci, M., Corea, G., Galeotti, F. & Lanzotti, V. The plant antifungal isoflavone genistein is metabolized by *Armillaria mellea* Vahl to give non-fungitoxic products. Plant Biosyst. - Int. J. Deal. Asp. Plant Biol. 140, 156–162 (2006).

70. Lah, L. et al. The versatility of the fungal cytochrome P450 monooxygenase system is instrumental in xenobiotic detoxification: Fungal P450 systems in xenobiotic detoxification. Mol. Microbiol. 81, 1374–1389 (2011).

71. Darwiche, R. et al. Plant pathogenesis–related proteins of the cacao fungal pathogen Moniliophthora perniciosa differ in their lipid-binding specificities. J. Biol. Chem. 292, 20558–20569 (2017).

72. Schneiter, R. & Di Pietro, A. The CAP protein superfamily: function in sterol export and fungal virulence. Biomol. Concepts 4, 519–525 (2013).

73. Gao, F. et al. Deacetylation of chitin oligomers increases virulence in soil-borne fungal pathogens. Nat. Plants 5, 1167–1176 (2019).

74. Mouyna, I. et al. What Are the Functions of Chitin Deacetylases in Aspergillus fumigatus? Front. Cell. Infect. Microbiol. 10, 28 (2020).

75. Saito, N. et al. Roles of RCN1, Regulatory A Subunit of Protein Phosphatase 2A, in Methyl Jasmonate Signaling and Signal Crosstalk between Methyl Jasmonate and Abscisic Acid. Plant Cell Physiol. 49, 1396–1401 (2008).

76. Cui, F., Brosché, M., Sipari, N., Tang, S. & Overmyer, K. Regulation of ABA dependent wound induced spreading cell death by MYB 108. New Phytol. 200, 634–640 (2013).

77. Mandaokar, A. & Browse, J. MYB108 Acts Together with MYB24 to Regulate Jasmonate-Mediated Stamen Maturation in Arabidopsis. Plant Physiol. 149, 851–862 (2009).

78. Liu, H. et al. Copper ions suppress abscisic acid biosynthesis to enhance defence against *Phytophthora infestans* in potato. Mol. Plant Pathol. 21, 636–651 (2020).

79. Maldonado, A. M., Doerner, P., Dixon, R. A., Lamb, C. J. & Cameron, R. K. A putative lipid transfer protein involved in systemic resistance signalling in Arabidopsis. Nature 419, 399–403 (2002).

80. O’Leary, B., Preston, G. M. & Sweetlove, L. J. Increased -Cyanoalanine β Nitrilase Activity Improves Cyanide Tolerance and Assimilation in Arabidopsis. Mol. Plant 7, 231–243 (2014).

81. Dodds, P. N. et al. Effectors of biotrophic fungi and oomycetes: pathogenicity factors and triggers of host resistance. New Phytol. 183, 993–1000 (2009).

82. Tanaka, S. & Kahmann, R. Cell wall–associated effectors of plant-colonizing fungi. Mycologia 113, 247–260 (2021).

83. Bass, A. J., Robinson, D. G. & Storey, J. D. Determining sufficient sequencing depth in RNA-Seq differential expression studies. http://biorxiv.org/lookup/doi/10.1101/635623 (2019) doi:10.1101/635623.

84. Urban, M., et al. PHI-base in 2022: a multi-species phenotype database for Pathogen–Host Interactions. Nucleic Acids Res. 50, D837–D847 (2022).

85. Have, A. ten, Mulder, W., Visser, J. & van Kan, J. A. L. The Endopolygalacturonase Gene *Bcpg1* Is Required for Full Virulence of *Botrytis cinerea*. Mol. Plant-Microbe Interactions® 11, 1009–1016 (1998).

86. Isshiki, A., Akimitsu, K., Yamamoto, M. & Yamamoto, H. Endopolygalacturonase Is Essential for Citrus Black Rot Caused by *Alternaria citri* but Not Brown Spot Caused by *Alternaria alternata*. Mol. Plant-Microbe Interactions® 14, 749–757 (2001).

87. Ökmen, B. et al. Detoxification of tomatine by *C ladosporium fulvum* is αL required for full virulence on tomato. New Phytol. 198, 1203–1214 (2013).

88. Ross-Davis, A. L. et al. Transcriptome of an *Armillaria* root disease pathogen reveals candidate genes involved in host substrate utilization at the host-pathogen interface. For. Pathol. 43, 468–477 (2013).

89. Ruiz-Dueñas, F. J. et al. Genomic Analysis Enlightens Agaricales Lifestyle Evolution and Increasing Peroxidase Diversity. Mol. Biol. Evol. 38, 1428–1446 (2021).

90. Koeck, M., Hardham, A. R. & Dodds, P. N. The role of effectors of biotrophic and hemibiotrophic fungi in infection: Effectors of biotrophic fungi. Cell. Microbiol. 13, 1849– 1857 (2011).

91. Flor, H. H. Current Status of the Gene-For-Gene Concept. Annu. Rev. Phytopathol. 9, 275–296 (1971).

92. Adelberg, J. et al. In vitro co-culture system for Prunus spp. and Armillaria mellea in phenolic foam rooting matric. Vitro Cell. Dev. Biol. - Plant 57, 387–397 (2021).

93. Ford, K. L., Baumgartner, K., Henricot, B., Bailey, A. M. & Foster, G. D. A native promoter and inclusion of an intron is necessary for efficient expression of GFP or mRFP in Armillaria mellea. Sci. Rep. 6, 29226 (2016).

94. Lam, K.-K., LaButti, K., Khalak, A. & Tse, D. FinisherSC: a repeat-aware tool for upgrading *de novo* assembly using long reads. Bioinformatics 31, 3207–3209 (2015).

95. Grabherr, M. G. et al. Full-length transcriptome assembly from RNA-Seq data without a reference genome. Nat. Biotechnol. 29, 644–652 (2011).

96. Grigoriev, I. V. et al. MycoCosm portal: gearing up for 1000 fungal genomes. Nucleic Acids Res. 42, D699–704 (2014).

97. Kuo, A., Bushnell, B. & Grigoriev, I. V. Fungal Genomics. in Advances in Botanical Research vol. 70 1–52 (Elsevier, 2014).

98. Jones, P. et al. InterProScan 5: genome-scale protein function classification. Bioinformatics 30, 1236–1240 (2014).

99. Almási, É. et al. Comparative genomics reveals unique wood decay strategies and fruiting body development in the Schizophyllaceae. New Phytol. 224, 902–915 (2019).

100. Emms, D. M. & Kelly, S. OrthoFinder: solving fundamental biases in whole genome comparisons dramatically improves orthogroup inference accuracy. Genome Biol. 16, (2015).

101. Emms, D. M. & Kelly, S. OrthoFinder: phylogenetic orthology inference for comparative genomics. Genome Biol. 20, 238 (2019).

102. Guy, L., Roat Kultima, J. & Andersson, S. G. E. genoPlotR: comparative gene and genome visualization in R. Bioinformatics 26, 2334–2335 (2010).

103. Katoh, K. & Standley, D. M. MAFFT multiple sequence alignment software version 7: improvements in performance and usability. Mol. Biol. Evol. 30, 772–780 (2013).

104. Capella-Gutierrez, S., Silla-Martinez, J. M. & Gabaldon, T. trimAl: a tool for automated alignment trimming in large-scale phylogenetic analyses. Bioinformatics 25, 1972–1973 (2009).

105. Stamatakis, A. RAxML version 8: a tool for phylogenetic analysis and post-analysis of large phylogenies. Bioinformatics 30, 1312–1313 (2014).

106. Nagy, L. G. et al. Latent homology and convergent regulatory evolution underlies the repeated emergence of yeasts. Nat. Commun. 5, 4471 (2014).

107. Lombard, V., Golaconda Ramulu, H., Drula, E., Coutinho, P. M. & Henrissat, B. The carbohydrate-active enzymes database (CAZy) in 2013. Nucleic Acids Res. 42, D490– D495 (2014).

108. Nguyen, L.-T., Schmidt, H. A., von Haeseler, A. & Minh, B. Q. IQ-TREE: A Fast and Effective Stochastic Algorithm for Estimating Maximum-Likelihood Phylogenies. Mol. Biol. Evol. 32, 268–274 (2015).

109. Revell, L. J. SIZE-CORRECTION AND PRINCIPAL COMPONENTS FOR INTERSPECIFIC COMPARATIVE STUDIES. Evolution 63, 3258–3268 (2009).

110. Revell, L. J. phytools: an R package for phylogenetic comparative biology (and other things): phytools: R package. Methods Ecol. Evol. 3, 217–223 (2012).

111. Steinegger, M. & Söding, J. MMseqs2 enables sensitive protein sequence searching for the analysis of massive data sets. Nat. Biotechnol. 35, 1026–1028 (2017).

112. Alexander, W. G., Wisecaver, J. H., Rokas, A. & Hittinger, C. T. Horizontally acquired genes in early-diverging pathogenic fungi enable the use of host nucleosides and nucleotides. Proc. Natl. Acad. Sci. 113, 4116–4121 (2016).

113. Wisecaver, J. H., Alexander, W. G., King, S. B., Todd Hittinger, C. & Rokas, A. Dynamic Evolution of Nitric Oxide Detoxifying Flavohemoglobins, a Family of Single-Protein Metabolic Modules in Bacteria and Eukaryotes. Mol. Biol. Evol. 33, 1979–1987 (2016).

114. Prospero, S., Holdenrieder, O. & Rigling, D. Comparison of the virulence of *Armillaria cepistipes* and *Armillaria ostoyae* on four Norway spruce provenances. For. Pathol. 34, 1–14 (2004).

115. Dobin, A. et al. STAR: ultrafast universal RNA-seq aligner. Bioinformatics 29, 15– 21 (2013).

116. Li, B. & Dewey, C. N. RSEM: accurate transcript quantification from RNA-Seq data with or without a reference genome. BMC Bioinformatics 12, 323 (2011).

117. Ritchie, M. E. et al. limma powers differential expression analyses for RNA-sequencing and microarray studies. Nucleic Acids Res. 43, e47–e47 (2015).

118. Robinson, M. D., McCarthy, D. J. & Smyth, G. K. edgeR: a Bioconductor package for differential expression analysis of digital gene expression data. Bioinforma. Oxf. Engl. 26, 139–140 (2010).

119. Plett, J. M. et al. Mycorrhizal effector PaMiSSP10b alters polyamine biosynthesis in *Eucalyptus* root cells and promotes root colonization. New Phytol. 228, 712–727 (2020).

