## Supplementary Note 1 for "Genomic innovation and horizontal gene transfer shaped plant colonization and biomass degradation strategies of a globally prevalent fungal pathogen"

**Extracellular virulence effectors identified from a conifer-specific stem-invasive pathosystem**

*In vitro* invasion assays using freshly cut conifer stems were set up to explore the differential gene expression profiles between higher and lower virulent isolates of two conifer-specific pathogenic species, *A. ostoyae* and *A. borealis* (Prospero *et al.* 2004; Heinzelmann *et al*. 2017)*.* We aimed to identify genes upregulated exclusively in the stem-invasive mycelia of the highly virulent isolates, presuming their contribution to more efficient invasive and necrotrophic activities. The search was confined to secretory proteins to determine extracellularly delivered potential plant interacting factors.

These experiments confirmed that the high-virulent A. *ostoyae* strain invaded the stems after 13 days, faster than the 19 days needed for the colonization by the less virulent one. The overall stem-invasive potential of the *A. borealis* isolates was slower as the virulent and less virulent isolates required an average of 23 and 24 days to colonize and showed a relatively small invasive advantage for the higher virulent strain.

For the comparative transcriptome analyses, we compared the stem-invasive transcriptomes of the highly virulent isolates (C18F / *A. ostoyae* and A6F / *A. borealis*) to those of the less virulent ones (C2F / *A. ostoyae* and A4F / *A. borealis*), as well as both highly virulent isolates to their respective vegetative mycelial controls (C18C / *A. ostoyae* and A6C / *A. borealis*). Finally, genes induced under plant invasive conditions but not in control vegetative mycelia were considered virulence-related genes. Among the pool of upregulated genes, we then determined the extracellularly targeted proteins and classified them as classically secreted proteins (CSPs) and leaderless secretory proteins (LSPs, identified as "extracellular" by using WolfPsort) (Agrawal *et al.* 2010).

In total, 443 and 232 genes from *A. borealis* and *A. ostoyae* were upregulated in the high-virulent isolates under plant invasive conditions. Following manual curation, from the *A. borealis* data, 73 inducible secretory proteins, including 23 SSPs (small secretory proteins, length < 300 amino acids), 15 genes with CAZy functions, five peptidases, two cerato-platanins, one intradiol ring-cleavage dioxygenase and two proteins with cupin1 domains could be identified. In the inducible secreted protein pool of *A. ostoyae*, 43 genes included 11 SSPs, 13 genes with CAZy functions and others representing two hydrophobins, two intradiol ring-cleavage dioxygenases and one tyrosinase were identified. In addition, four FAD-monooxygenases from *A. ostoyae* and four Cytochrome P450 genes from *A. borealis* were prevalent in their LSP-related gene pools.

Notably, 29 genes from the inducible extracellular pools of the two species were homologous to already identified pathogenicity or virulence factors (based on PHI-base; Urban *et al.* 2022). Amongst them, both species had pectinase (GH28) and cellulose-degrading (GH3) genes related to plant interacting effectors. GH28 genes have previously been recognized as encoding virulence effectors in several ascomycete pathogens; namely in *Claviceps purpurea*, a grass and cereal pathogen (Oeser et al., 2002), and *Alternaria citri*, responsible for black rot of citrus fruits (Isshiki et al., 2001), and as a crucial factor for reaching the full virulence potential in *Botrytis cinerea* (Have et al., 1998). Regarding GH3, earlier studies have linked GH3 to the "tomatinase" function involved in removing the terminal β-1,2-d-glucose and β-1,3-d-xylose residues from the antifungal glycoalkaloid α-tomatine. As a result, GH3 enzymes can play an essential role in overcoming plant defense (Martin-Hernandez et al., 2002; Ökmen et al., 2013). Both of these glycoside hydrolase families were also found significantly upregulated in the initial stage of infection by other necrotrophic white rot pathogens *Heterobasidion occidentale* and *H. irregulare* when infecting Norway spruce (Hu *et al,* 2020).

In addition, members of other glycoside hydrolase families, single GH15 and GH7 genes were identified from the *A. ostoyae* pool*,* and two GH51 genes and one GH131 gene were upregulated in *A. borealis.* Interestingly, the gene with the GH15 domain also contained a CBM20 domain, both previously reported to be required for *B. cinerea* necrosis (Yang et al., 2018). Similarly, a GH7 homologue, regarded as a crucial virulence factor in *Phytophthora sojae,* has also been described as a cell death-triggering factor in different hosts (Tan et al., 2020). In contrast to necrosis factors, the secreted GH131 β-glucanases are widespread among all pathogenic, saprobic and symbiotic fungi and have been found essential for colonising plant tissues (Anasontzis et al., 2019).

Regarding other CAZy functions, two multicopper oxidases (AA1) were also present in the virulent pool of *A. ostoyae,* and they were homologous to the BcLCC2 laccase of *B. cinerea*, considered as a factor affecting virulence by being part of a laccase-mediated system involved in detoxifying a broad-spectrum of antibiotic, possibly aromatic compounds (Schouten *et al, 2002, 2008*).

Considering non-CAZymes, palmitoyl protein thioesterase (Ppt1) from the *A. borealis* pool was homologous to the Ppt1 gene, regarded as a virulence factor, from *Cochliobolus heterostrophus* and *C. miyabeanus* (Zainudin *et al*, 2015). Other possible virulence factors present in the *A. borealis* pool were a carboxylesterase B gene, homologous to a *Fusarium graminearum* virulence factor (Zhang *et al,* 2016), and two cerato-platanins, corresponding to cell death-inducing proteins involved in the pathogenicity of *Heterobasidion annosum* and *Sclerotinia sclerotiorum* (Chen *et al*, 2015; Pan *et al,* 2018). Furthermore, a gene related to subtilisin-like proteases from *A. borealis* contained an inhibitory I9 and a peptidase S8 domain, the same as its homolog Bcser2 from *B. cinerea* where it has been confirmed as crucial for the virulence, conidiation and sclerotial formation (Liu *et al.*, 2020).

Among the remaining CSPs in both species, intradiol ring-cleavage dioxygenase and sodium dismutase (Cu/Zn) genes were the most notable. The dioxygenases are homologous to the pcaH gene, involved in the 4-hydroxybenzoate degradation pathway, which is vital for the pathogenicity of Xanthomonas campestris (Wang et al., 2015). The superoxide dismutases (SOD) may contribute to the invasive necrotrophic activities as the deletion of the Bcsod1 gene in *Botrytis* *cinerea* displayed reduced virulence (López-Cruz et al. 2017).

Finally, amongst the LSPs of *A. ostoyae*, we identified four FAD-monooxygenases related to the *MAK1* genes of *Fusarium solani* involved in degrading plant phytoalexins (Enkerli *et al*, 1998). An additional lactonase gene was homologous to the 3-carboxy-cis,cis-muconate lactonizing enzyme (CMLE) in *Fusarium oxysporum* whose deletion, in addition to the loss of pathogenicity, resulted in an inability to catabolize phenolic compounds via the β-ketoadipate pathway (Michielse *et al., 2012*).

We used Orthogroup membership to determine species-specific or shared potential virulence factors between *A. ostoyae* and *A. borealis.* Among the classically secreted proteins, orthogroups OG0000124 (intradiol ring-cleavage dioxygenase), OG0000002 (GMC oxidoreductase), OG0000817 (Kre9/Knh1 family), OG0001012 (Cu/Zn superoxide dismutase) and OG0004292 (GH28) were shared between the two species.

Regarding the species-specific orthogroups, in the *A. ostoyae* pool, hydrophobins, multicopper oxidases and GH28 genes, while in the *A. borealis* pool, cerato-platanins, S10 peptidases, alpha-arabinofuranosidases, and proteins with either cupin1 or LysM domains were identified with identical orthogroup IDs representing each group only with two genes.

Amongst the LSPs, *A. ostoyae* had four FAD-monooxygenases with the same orthogroup ID (OG0000058) in addition to one cytochrome P450 orthogroup (OG0001710) that was shared between both species.

**References**

Heinzelmann, R., Prospero, S., and Rigling, D. (2017). Virulence and Stump Colonization Ability of *Armillaria borealis* on Norway Spruce Seedlings in Comparison to Sympatric *Armillaria* Species. Plant Dis. *101*, 470–479.<https://doi.org/10.1094/PDIS-06-16-0933-RE>.

Prospero, S., Holdenrieder, O., and Rigling, D. (2004). Comparison of the virulence of *Armillaria cepistipes* and *Armillaria ostoyae* on four Norway spruce provenances. For. Pathol. *34*, 1–14.<https://doi.org/10.1046/j.1437-4781.2003.00339.x>.

Agrawal, G.,K., Jwa, NS., Lebrun, MH., Job, D., Rakwal, R. (2010). Plant secretome: Unlocking secrets of the secreted proteins. Proteomics, 10, 799-827. https://doi.org/10.1002/pmic.200900514

Urban, M., Cuzick, A., Seager, J., Wood, V., Rutherford, K., Venkatesh, S.Y., Sahu, J., Iyer, S.V., Khamari, L., De Silva, N., et al. (2022). PHI-base in 2022: a multi-species phenotype database for Pathogen–Host Interactions. Nucleic Acids Res. 50, D837–D847.<https://doi.org/10.1093/nar/gkab1037>.

Oeser, B., Heidrich, P. M., Müller, U., Tudzynski, P., & Tenberge, K. B. (2002). Polygalacturonase is a pathogenicity factor in the *Claviceps purpurea*/rye interaction. Fungal Genet. Biol, 36(3), 176-186.<https://doi.org/10.1016/S1087-1845(02)00020-8>.

Isshiki, A., Akimitsu, K., Yamamoto, M., & Yamamoto, H. (2001). Endopolygalacturonase is essential for citrus black rot caused by *Alternaria citri* but not brown spot caused by *Alternaria alternata.* Mol. Plant Microbe Interact, 14(6), 749-757. https://doi.org/10.1094/MPMI.2001.14.6.749

Have, A. T., Mulder, W., Visser, J., & van Kan, J. A. (1998). The endopolygalacturonase gene Bcpg1 is required for full virulence of *Botrytis cinerea*. Mol. Plant Microbe Interact, *11*(10), 1009-1016.<https://doi.org/10.1094/MPMI.1998.11.10.1009>

Martin-Hernandez, A. M., Dufresne, M., Hugouvieux, V., Melton, R., & Osbourn, A. (2000). Effects of targeted replacement of the tomatinase gene on the interaction of *Septoria lycopersici* with tomato plants. Mol. Plant Microbe Interact, 13(12), 1301-1311. https://doi.org/10.1094/MPMI.2000.13.12.1301

Ökmen, B., Etalo, D. W., Joosten, M. H., Bouwmeester, H. J., de Vos, R. C., Collemare, J., & de Wit, P. J. (2013). Detoxification of α‐tomatine by *Cladosporium fulvum* is required for full virulence on tomato. New Phytol., 198(4), 1203-1214. <https://doi.org/10.1111/nph.12208>

Hu, Y., Elfstrand, M., Stenlid, J., Durling, M. B., & Olson, Å. (2020). The conifer root rot pathogens *Heterobasidion irregulare* and *Heterobasidion occidentale* employ different strategies to infect Norway spruce. Sci Rep, 10(1), 1-10.<https://doi.org/10.1038/s41598-020-62521-x>

Yang, C., Liang, Y., Qiu, D., Zeng, H., Yuan, J., & Yang, X. (2018). Lignin metabolism involves *Botrytis cinerea* BcGs1-induced defense response in tomato. BMC Plant Biol., *18*(1), 1-15.<https://doi.org/10.1186/s12870-018-1319-0>

Tan, X., Hu, Y., Jia, Y., Hou, X., Xu, Q., Han, C., & Wang, Q. (2020). A conserved glycoside hydrolase family 7 cellobiohydrolase PsGH7a of *Phytophthora sojae* is required for full virulence on soybean. Front. Microbiol. 11, 1285.<https://doi.org/10.3389/fmicb.2020.01285>

Anasontzis, G. E., Lebrun, M. H., Haon, M., Champion, C., Kohler, A., Lenfant, N., Martin, F., O'Connell, R.J., Riley, R., Grigoriev, I.V., Henrissat, B., Berrin, J.-G., & Rosso, M. N. (2019). Broad‐specificity GH131 β‐glucanases are a hallmark of fungi and oomycetes that colonize plants. Environ. Microbiol. *21*(8), 2724-2739.<https://doi.org/10.1111/1462-2920.14596>

Schouten, A., Wagemakers, L., Stefanato, F. L., Kaaij, R. M. V. D., & Kan, J. A. V. (2002). Resveratrol acts as a natural profungicide and induces self‐intoxication by a specific laccase. Mol. Microbiol*.*, *43*(4), 883-894.<https://doi.org/10.1046/j.1365-2958.2002.02801.x>

Schouten, A., Maksimova, O., Cuesta‐Arenas, Y., Van Den Berg, G., & Raaijmakers, J. M. (2008). Involvement of the ABC transporter BcAtrB and the laccase BcLCC2 in defence of *Botrytis cinerea* against the broad‐spectrum antibiotic 2, 4‐diacetylphloroglucinol. Environ. Microbiol*.*, *10*(5), 1145-1157.<https://doi.org/10.1111/j.1462-2920.2007.01531.x>

Liu, X., Xie, J., Fu, Y., Jiang, D., Chen, T., & Cheng, J. (2020). The subtilisin-like protease Bcser2 affects the sclerotial formation, conidiation and virulence of *Botrytis cinerea*. *Int. J.* Mol. Sci., *21*(2), 603.<https://doi.org/10.3390/ijms21020603>

Zainudin, N. A. I. M., Condon, B., De Bruyne, L., Van Poucke, C., Bi, Q., Li, W., Höfte, M. & Turgeon, B. G. (2015). Virulence, host-selective toxin production, and development of three *Cochliobolus phytopathogens* lacking the Sfp-type 4′-phosphopantetheinyl transferase Ppt1. Mol. Plant Microbe Interact., 28(10), 1130-1141.<https://doi.org/10.1094/MPMI-03-15-0068-R>

Zhang, Y., He, J., Jia, L. J., Yuan, T. L., Zhang, D., Guo, Y., Wang, Y., & Tang, W. H. (2016). Cellular tracking and gene profiling of *Fusarium graminearum* during maize stalk rot disease development elucidates its strategies in confronting phosphorus limitation in the host apoplast. PLoS Pathog*.*, *12*(3), e1005485.<https://doi.org/10.1371/journal.ppat.1005485>

Chen, H., Quintana, J., Kovalchuk, A., Ubhayasekera, W., & Asiegbu, F. O. (2015). A cerato-platanin-like protein HaCPL2 from *Heterobasidion annosum sensu stricto* induces cell death in *Nicotiana tabacum* and *Pinus sylvestris*. Fungal Genet. Biol, *84*, 41-51.<https://doi.org/10.1016/j.fgb.2015.09.007>

Pan, Y., Wei, J., Yao, C., Reng, H., & Gao, Z. (2018). SsSm1, a Cerato-platanin family protein, is involved in the hyphal development and pathogenic process of *Sclerotinia sclerotiorum*. Plant Sci., *270*, 37-46.<https://doi.org/10.1016/j.plantsci.2018.02.001>

Wang, J. Y., Zhou, L., Chen, B., Sun, S., Zhang, W., Li, M., Tang, H., Jiang, B. L., Tang, J. L. & He, Y. W. (2015). A functional 4-hydroxybenzoate degradation pathway in the phytopathogen *Xanthomonas campestris* is required for full pathogenicity. Sci. Rep., *5*(1), 1-13.<http://doi.org/10.1038/srep18456>

López-Cruz, J., Óscar, C. S., Emma, F. C., Pilar, G. A., & Carmen, G. B. (2017). Absence of Cu–Zn superoxide dismutase BCSOD1 reduces *Botrytis cinerea* virulence in Arabidopsis and tomato plants, revealing interplay among reactive oxygen species, callose and signalling pathways. Mol. Plant Pathol., *18*(1), 16-31.<https://doi.org/10.1111/mpp.12370>

Enkerli, J., Bhatt, G., & Covert, S. F. (1998). Maackiain detoxification contributes to the virulence of *Nectria haematococca* MP VI on chickpea. Mol. Plant Microbe Interact., *11*(4), 317-326. <https://doi.org/10.1094/MPMI.1998.11.4.317>

Michielse, C. B., Reijnen, L., Olivain, C., Alabouvette, C., & Rep, M. (2012). Degradation of aromatic compounds through the β‐ketoadipate pathway is required for pathogenicity of the tomato wilt pathogen *Fusarium oxysporum f. sp. lycopersici*. Mol. Plant Pathol., *13*(9), 1089-1100.<https://doi.org/10.1111/j.1364-3703.2012.00818.x>

Siewers, V., Viaud, M., Jimenez-Teja, D., Collado, I. G., Gronover, C. S., Pradier, J. M., Tudzynsk, B., & Tudzynski, P. (2005). Functional analysis of the cytochrome P450 monooxygenase gene bcbot1 of *Botrytis cinerea* indicates that botrydial is a strain-specific virulence factor. Mol. Plant Microbe Interact., *18*(6), 602-612.<https://doi.org/10.1094/MPMI-18-0602>
