## Supplementary for "Genomic innovation and horizontal gene transfer shaped plant colonization and biomass degradation strategies of a globally prevalent fungal pathogen"

**This file contains Figure legends for:**

**Figure S1**

**Figure S2**

**Figure S3**

**Figure S4**

**Figure S5**

**Figure S6**

**Figure S7**

**Figure S8**

**Figure S9**

**Figure S10**

**Figure S11**

**Figure S12**

**Figure S13**

**Figure S14**

**Figure S15**

**Figure S16**

**Figure S17**

**Figure S18**

**Figure S19**

**Figure S20**

**Figure S21**

**Figure S22**

**Other supplementary files:**

**Table S1.** New Armillaria genomes, list of species in each dataset used in this study, and their respective species trees

**Table S2.** Enriched genes in Armillaria duplications, and novel-core genes in Armillaria clade.

**Table S3.** Carbohydrate-active enzymes (CAZymes) and plant cell wall

degrading enzymes (PCWDEs) identified in Dataset2, copy numbers of substrate-based PCWDEs in each species, PCA Loadings from phylogenetic PCA, co-enriched CAZy orthogroups and their domain architecture

**Table S4.** Horizontal gene transfers in Physalacriaceae using ailen-index and phylogenetic validation form gene trees

**Table S5.** Differentially expressed genes in 6 RNA seq experiments and odds ratio for different functional categories

**Table S6.** Expression data for *A. luteobubalina* small secreted proteins in the *in-planta* assay, and virulence factors and orthogroup counts in stem-invasion assays.

**Figure Legends**

**Figure S1: Summary of gene losses and gains from COMPARE mappings.** Duplications (green) and losses (red) at each node for Dataset1. Bootstrap support values less than 80 are shown in blue.

**Figure S2: GO terms enriched in duplicated genes.** Significantly enriched GO terms in the 1473 orthogroups, inferred by 2913 duplications at *Armillaria* MRCA. X-axis shows the percentage of significant genes from the total number of genes, y-axis shows p-values. Blue shows lower and red shows higher p-values. GO terms that had at least 30% of genes significant from the total number of genes are mentioned on the plot (see Table S2 for the complete list of enriched GO terms)

**Figure S3: Novel-core genes in Armillaria.** UpsetR plot showing novel-core orthogroups present exclusively in at least 12 *Armillaria* species (including Guynagaster) (for list of Orthogroups and their functional InterPro annotations, see Table S2)

**Figure S4: Synteny of the bioluminescence cluster in *Armillaria*.** Syntenic regions of the 5 bioluminescence genes in *Armillaria*

**Figure S5: Plant biomass degradation genes in *Armillaria:*** Phylogenetic PCAs and their respective loading factors for PCWDE gene families. Species abbreviations are colored according to nutritional modes.

**Figure S6: Barplots for Hemicellulase genes.** Gene copy numbers of hemicellulose-acting CAZymes. Bars are colored according to nutritional mode. X-axis shows gene copy numbers and Y-axis depicts species from selected lifestyles (Physac, LD and WR) sorted according to their phylogenetic tree as in Dataset2. Dashed vertical line depicts average gene copies for each family (for complete list of gene copy numbers in all species of Dataset2 - Table S3).

**Figure S7: Barplots for Ligninase genes.** Gene copy numbers of lignin-acting CAZymes. Bars are colored according to nutritional mode. X-axis shows gene copy numbers and Y-axis depicts species from selected lifestyles (Physac, LD and WR) sorted according to their phylogenetic tree as in Dataset2. Dashed vertical line depicts average gene copies for each family (for complete list of gene copy numbers in all species of Dataset2 - Table S3).

**Figure S8: Barplots for Putative ligninase genes.** Gene copy numbers of CAZymes putatively acting on lignin monomers. Bars are colored according to nutritional mode. X-axis shows gene copy numbers and Y-axis depicts species from selected lifestyles (Physac, LD and WR) sorted according to their phylogenetic tree as in Dataset2. Dashed vertical line depicts average gene copies for each family (for complete list of gene copy numbers in all species of Dataset2 - Table S3).

**Figure S9: Barplots for Cellulase genes.** Gene copy numbers of cellulose-acting CAZymes. Bars are colored according to nutritional mode. X-axis shows gene copy numbers and Y-axis depicts species from selected lifestyles (Physac, LD and WR) sorted according to their phylogenetic tree as in Dataset2. Dashed vertical line depicts average gene copies for each family (for complete list of gene copy numbers in all species of Dataset2 - Table S3).

**Figure S10: Barplots for Pectinase genes.** Gene copy numbers of pectin-acting CAZymes. Bars are colored according to nutritional mode. X-axis shows gene copy numbers and Y-axis depicts species from selected lifestyles (Physac, LD and WR) sorted according to their phylogenetic tree as in Dataset2. Dashed vertical line depicts average gene copies for each family. Dashed vertical line depicts average gene copies for each family (for complete list of gene copy numbers in all species of Dataset2 - Table S3).

**Figure S11: Expression of HT and VT genes in *A. ostoyae* developmental transcriptome.** Violin plot showing gene expression of phylogenetically validated HT and VT genes in *A. ostoyae* fruiting body development transcriptome. Y-axis shows log_2_ transformed expression values, and x-axis shows the sample comparisons for each experiment.

**Figure S12: Experimental setup.** Setup for the new RNA Seq experiments used in this study. A) Setup for the time-course experiment. B) Setup for the stem invasion assay.

**Figure S13: Enrichment of differentially expressed genes of selected gene families in 6 RNA-Seq datasets.** The heatmap shows enrichment ratios for 23 gene groups (“Ergothione: removed due to no enrichment) from aggregated differential gene expression data across 6 experiments (A - upregulated, B - downregulated genes). Y-axis shows the sample comparison for each dataset, with number of DEGs shown as a barplot at right. In the heatmap, warmer colors mean higher enrichment ratios (for complete list of odds ratios, see Table S5).

**Figure S14: Expression heatmaps in *A. luteobubalina*.** Heatmaps showing gene expression along the time course in *A. luteobubalina* for gene families related to host immune suppression, oxidative stress, detoxification, and cytotoxicity. Warmer color depicts higher expression.

**Figure S15: Transcriptomic analysis in *A.luteobubalina* and E.*grandis*.** A) Multidimensional scaling showing grouping of biological replicates for *A. luteobubalina* (left) and *E. grandis* (right). Replicates from each time point are shown in different color. B) Barplot of upregulated (purple) and downregulated genes (yellow) in *A. luteobubalina* (left) and *E. grandis* (right).

**Figure S16: Validation of Transcriptomics data by SuperSeq.** Percentage of differentially expressed genes (y-axis) obtained at current (dashed vertical line) and estimated proportion read depth (x-axis) for different sample comparisons in in-planta and stem-invasion assays. Colors represent different sample comparisons.

**Figure S17: GO terms enriched in five significant STEM profiles in *A. luteobubalina*.** Line graph in blue shows expression trend across the time series, bold line depicts the average expression trend of the STEM profile. Dotplots are colored according to their significant p-values, with blue for high and red for low. Dot sizes are proportional to number of significant genes enriched from the total number of genes for each GO term.

**Figure S18 GO terms enriched in four significant STEM profiles in *E. grandis*.** Line graph in green shows expression trend across the time series, bold line depicts the average expression trend of the STEM profile. Dotplots are colored according to their significant p-values, with blue for high and red for low. Dot sizes are proportional to number of significant genes enriched from the total number of genes for each GO term.

**Figure S19: Expression of Differentially expressed genes (DEGs) in *E. grandis*.** A) Heatmap of all DEGs across the timecourse; B) Venn diagram of common and unique upregulated genes; C) Venn diagram of common and unique downregulated genes

**Figure S20: Pathogenicity-induced SSP expression and conservation in *A. luteobubalina*.** A) Heatmap shows log_2_ fold changes for annotated SSPs, upregulated in at least one-time point. Red shows higher and blue depicts lower logFC; followed by presence/absence matrix of homologs in 131 species (Dataset 2). X-axis shows ProteinIDs for both heatmap and presence/absence matrix. Y-axis shows sample comparisons in the heatmap; and species order in presence/absence matrix.

**Figure S21: Copy numbers and conservation of Armlut1_1348401.** A) Summary of copy-numbers of the orthogroup OG0000784, comprising PiSSP Armlut1_1348401. B) Trimmed multiple sequence alignment of proteins in OG0000784.

**Figure S22: Copy numbers and conservation of Armlut1_1165297.** A) Summary of copy-numbers of the orthogroup OG0000401, comprising PiSSP Armlut1_1165297. B) Trimmed multiple sequence alignment of proteins in OG0000401.
